# Genome-wide signatures of geographic expansion and breeding process in soybean

**DOI:** 10.1101/2022.03.24.485586

**Authors:** Ying-hui Li, Chao Qin, Li Wang, Chengzhi Jiao, Huilong Hong, Yu Tian, Yanfei Li, Guangnan Xing, Jun Wang, Yongzhe Gu, Xingpeng Gao, Delin Li, Hongyu Li, Zhangxiong Liu, Xin Jing, Beibei Feng, Tao Zhao, Rongxia Guan, Yong Guo, Jun Liu, Zhe Yan, Lijuan Zhang, Tianli Ge, Xiangkong Li, Xiaobo Wang, Hongmei Qiu, Wanhai Zhang, Xiaoyan Luan, Yingpeng Han, Dezhi Han, Ruzhen Chang, Yalong Guo, Jochen C. Reif, Scott A. Jackson, Bin Liu, Shilin Tian, Li-juan Qiu

## Abstract

The clarification of genomic signatures left during evolutionary histories of crops is crucial for breeding varieties adapting to changing climate. Soybean, a leguminous crop, provides both plant oil and protein. Here, we analyzed genome sequences of 2,214 soybeans and proposed its evolutionary route, which includes four geographic paths, expansion of annual wild soybean (*Glycine soja* Sieb. & Zucc.) from Southern China, domestication in Central China, expansion of landrace (*G. max* (L.) Merr.), and local breeding. We observed that local adaptation of the wild and cultivated soybeans was largely independent, and that genetic introgression was mostly derived from sympatric rather than allopatric wild populations during the range expansion of soybean landraces. Range expansion and breeding processes were accompanied with positive selection of flowering-time genes including *GmSPA3c* as validated by knock-out mutants. Our study shed lights on the evolutionary history of soybean and provides valuable genetic resources for future breeding.

**Teaser:** The expansion and selection history of soybean

## Introduction

Plants evolution is an expansive process that includes early domestication, habitat expansion and subsequent genetic improvement (*1*), reflecting the impact of artificial and natural selection on gene diversity. During evolution, genes favoring intensive cultivation, high productivity and quality were selected and resulted in the appearance of landraces and subsequent improved cultivars. The spread of landraces and improved cultivars led to substantial increases in range of adaptation and productivity. During habitat expansion, in contrast, gene variants accumulate, which allowed adaptation to new environmental conditions. Therefore, a retrospective view of changes in genetic diversity can be used to identify genes that are crucial, for example, to the future adaptation of crops to changing climate.

Soybean is a remarkable crop with rich genomic resources (*2*), a worldwide leading source of protein and oils, including edible oil, human food, livestock forage, and biodiesel. Cultivated soybean (*Glycine max* (L.) Merr.) was proposed to have been domesticated in China about 5,000 years ago from its annual wild relative (*Glycine soja* Sieb. & Zucc.)(*3*). After domestication, local landraces spread throughout East Asia, sympatric, i.e. sharing the habitat, with their wild relatives. The success of modern breeding led to the replacement of local landraces by high-yielding and quality cultivars in the soybean production. Soybean is an excellent system to study how demography and selection altered crop genomes. The molecular footprint left during domestication and genetic improvement had been clarified in several studies (*2, 4-6*), however, little is known the landscape of genomic signatures underlying the expansion of the wild soybean and landrace respectively, and if the gene flow between sympatric wild soybean and landraces facilitated the local adaptation, the same as maize, sorghum etc. (*7, 8*).

With newly sequenced 1,674 soybean genomes and 540 previously released genomes (*4, 9*) covering its geographic distribution and the diversity of cultivars, we clarified the spreading routes of soybean, examined the prevalence of introgression from the wild to cultivated populations, detected signatures of selection in different evolutionary processes and validated the function of one representative flowering time gene involving the expansion of cultivated soybean with the CRISPR/Cas9 knock-out experiments.

## Results

### A genomic variation map of soybean

The subgenus *Soja* includes domesticated *G. max* and its wild antecedent, *G. soja*. The *G. max* species includes landraces and improved cultivars. To analyze genetic variation, an extensive and diverse set of soybean genomes was studied, including 1,993 *G. max* (1,131 landraces and 862 improved cultivars), 218 *G. soja*, two perennial *G. tomentella* and one perennial *G. tabacine* (**Figs. 1A, 1B and table S1**). Of these, 1,674 genomes were newly sequenced and 540 were previously published(*4, 9*). *G. soja* accessions and *G. max* landraces were collected from their native geographic range, i.e. East Asia. Improved cultivars were sampled globally, mainly from primary soybean producing countries such as the United States of America, Japan, Korea, and China (**table S1**). A total of 1,690 out of the 1,993 cultivated soybeans (84.8%), were selected from the Chinese primary and applied core collections based on 14 agronomic traits and sequences of 60 single copy loci, representing the broad genetic diversity of the 23,587 cultivated soybeans from the Chinese National Soybean Gene Bank(*10, 11*).

**Fig. 1.**
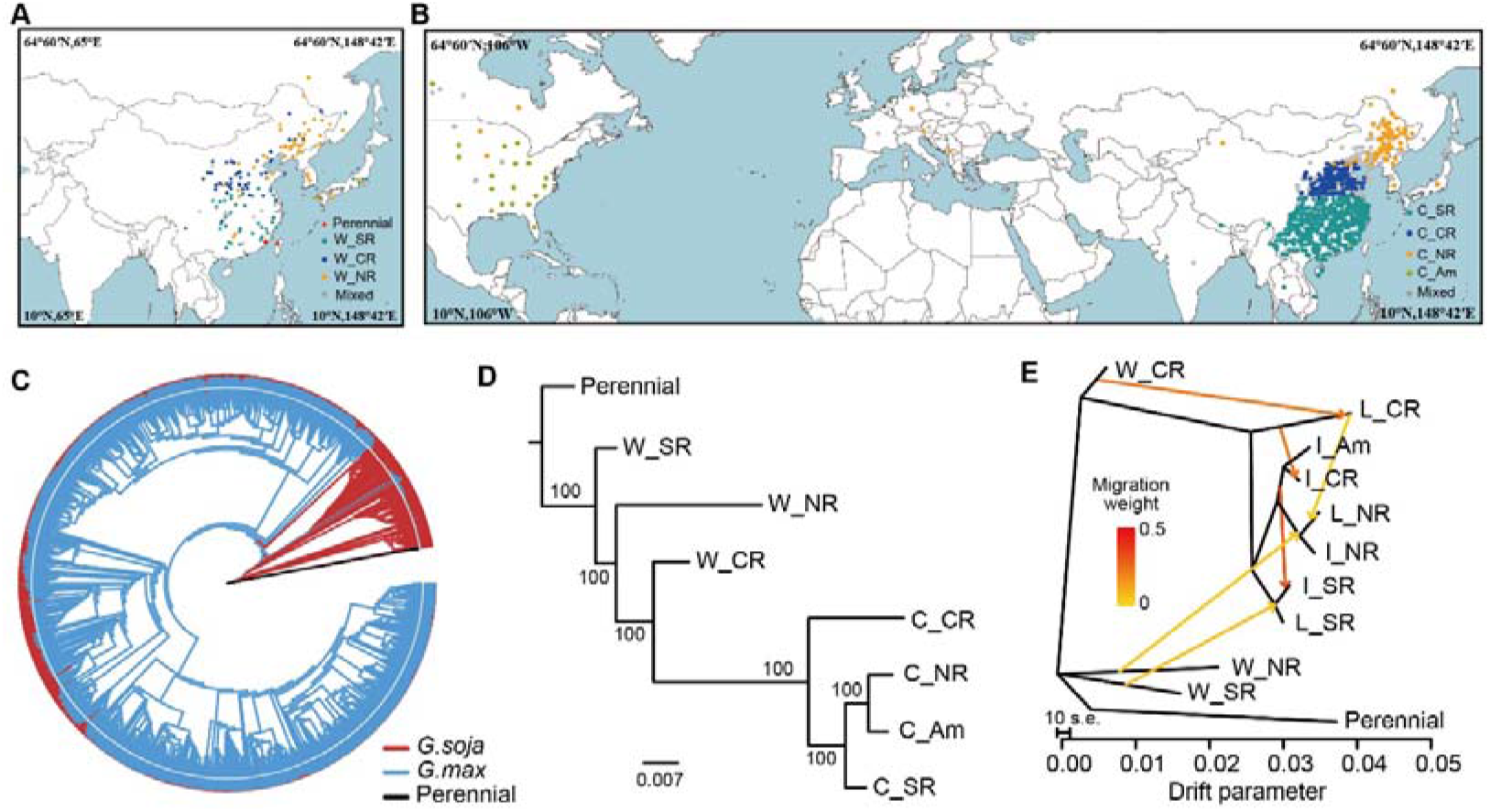
The distribution, genetic diversity and population structure of Subgenus *Soja*. (**A**) Geographic distribution of *G. soja* accessions. (**B**) Geographic distribution of *G. max* accessions. (**C**) Population structure of 1,993 *G. max* and 218 *G. soja* accessions using three perennial accessions as the outgroup. The outer ring indicated the estimated proportions of an individual’s assignment at *K* = 2. (**D**) Trees for seven subpopulations inferred from population structure analysis with three perennial accessions as the outgroup. The percentage bootstrap support is indicated at each node. (**E**) TreeMix analysis of soybean groups with three perennial accessions as the outgroup and m = 6. The arrow indicated the migration direction. Abbreviations: “W_” represented the wild soybean, “C_” denoted the cultivated soybean, “L_” represented the landraces and “I_” denoted the improved cultivars; SR indicated the Chinese Southern region; CR implied the Chinese Central region surrounding the mid-down stream of Yellow River valley; NR stand for the Chinese Northern region plus Japan, Korean peninsula and Russian Far East region.

A total of 16.41 Tb (Tera bases) high-quality genome sequences of 2,214 accessions were mapped to the soybean reference genome(*12*) (**table S1**). We obtained 8,785,134 high-confidence biallelic SNPs (Supplementary Methods), and subsequently annotated 1,259,917 SNPs (14.3%) located within 53,720 protein-coding genes (95.85% of the total genes) (**fig. S1, 2 and table S2**). The validation ratio of 98.3% for 114 accessions randomly selected from all 2,214 samples using the genotyping by target sequencing method, indicated that SNP calling in this study had a low miscall rate (**table S3**). Not surprisingly, SNP density was significantly higher in promoter regions than in coding regions (p ≅ 0; **fig. S3**). A total of 170,193 missense and 5,214 stop-gain/loss SNPs were observed that caused amino acid changes, premature stops or elongated transcripts, respectively, leading to potential changes in 40,742 protein sequences (72.70% of the total) (**table S4**).

### Wild and cultivated soybeans showed an analogous population differentiation according to their geographic origin

To further elucidate the genetic structure in soybeans, we selected 1,721,062 SNPs with weak linkage disequilibrium (LD; *r*^2^ < 0.8) amongst each other(*13*). Three methods were used to infer population structure including the neighbor-joining (NJ) tree, principal components (PCA)(*14*) and Bayesian clustering(*15*). The results from each were concordant (**Fig. 1c and fig. S4-5**) and showed that the primary genetic differentiation was between *G. max* and its progenitor *G. soja*, indicating that soybean underwent a single domestication event consistent with previous studies(*4, 16*). We subsequently found that 204 *G. soja* accessions (93.6%) were clearly distinguished into three sub-populations corresponding to three different geographical regions: (1) the Chinese Northern region plus Japan, Korean peninsula, and Russian Far East region (termed as W_NR, n=89, “W” indicated “wild”); (2) Chinese Central region surrounding the mid-down stream of the Yellow River valley (W_CR, n=73); and (3) the Chinese Southern region (W_SR, n=42) (**fig. S4A, 5B, and 6A**). Genotypes of the W_SR formed the first divergent clade in the maximum likelihood rooted tree, indicating that the wild soybean likely spread northward from the Chinese Southern region (**Fig. 1C**). Similarly, 1,632 *G. max* accessions (81.9%) were distinguished into four different geographical regions, i.e., C_NR (n=403, “C” indicated “cultivated”), C_CR (n=278), C_SR (n=711), and the America (C_Am, n=240) (**fig. S4C, 5C, 6B and 7**), demonstrating an analogous population differentiation with its sympatric wild relatives. To gain insights into the origin locations of the cultivated soybean, we constructed a neighbor-joining tree of all pooled sub-populations with perennial species as outgroup **(Fig. 1D)**. It revealed that the wild subpopulation from the central region (W_CR) is phylogenetically closest to the cultivated clade and the landraces from the central region (L_CR) is the first subclade diverged from the wild subpopulations. Meanwhile, the estimate of effective population sizes (*N*_e_) in the three sub-populations of cultivated soybeans revealed that C_SR and C_NR showed stronger bottlenecks than C_CR (**fig. S8**), suggesting that the cultivated populations were expanded from C_CR to C_SR and C_NR. Furthermore, we detected gene flow from the Central to Northern landrace populations and from the Central to the Southern improved cultivars (**Fig. 1E**). Taken together, these results implied that the middle reaches of the Yellow River as the domestication center of soybeans.

### Introgressions facilitated local adaptation of landraces

Soybean was initially constrained and adapted to a narrow and specific geographic range(*3*), but subsequently underwent a massive spread after its domestication in Central China. One questions is whether gene flow from local wild populations facilitated its adaptation and spread, as reported for other crop species(*7, 17*). In order to answer this question, we first inferred the gene flow between inter-/intra-sub-populations of *G. soja* and *G. max* at the genome-wide level using TreeMix(*18*). Noticeably, we observed directional gene flow from local wild to landrace populations among all three subpopulations (**Fig. 1E and fig. S9**). Next, we calculated *f*_d_ values in 10 kb non-overlapping sliding windows to define the genomic regions of gene flow. We found that the *f*_d_ values were significantly higher for comparisons that involved two sympatric wild and landrace populations than that those involving allopatric populations (*p* < 2.97e-10, *t* test) (**Fig. 2**). This was true except for introgression levels from the Southern wild to the Southern landrace populations which was not different from the Northern wild to the Southern landrace populations (**Fig. 2**; *p* = 0.1, *t* test), which might reflect the genetic exchange between the wild populations during the last glacier maximum(*19*). These data indicate that when soybean landraces migrated to the Southern and Northern regions, the gene flow from local wild populations likely accelerated local adaptation.

**Fig. 2.**
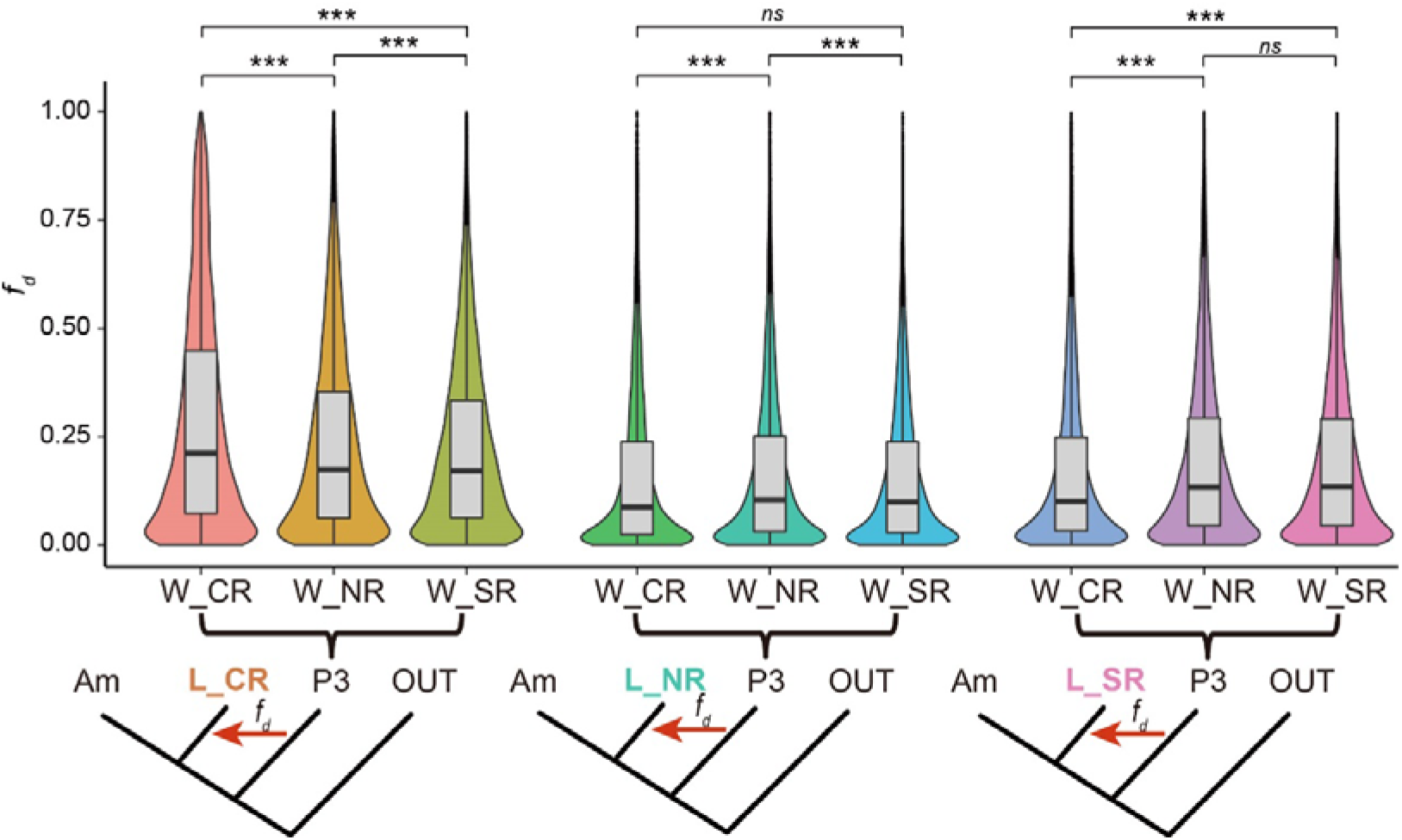
The *f*_d_ values were calculated for each landrace sub-population (denoted as the colored population on the tree) with potential introgression from three wild sub-populations, respectively. The abbreviation “W_” indicates the wild soybean; “L_” represents the landraces. “*ns*” means “not significant” (*p* > 0.05). SR indicated the Chinese Southern region; CR implied the Chinese Central region surrounding the mid-down stream of Yellow River valley; NR stood for the Chinese Northern region plus Japan, Korean peninsula and Russian Far East region. “Am” pointed to America.

We had a closer look on windows including the top 5% *f*_d_ values and functionally annotated genes associated with seed quality, flowering time, and biotic resistance **(table S5; fig. S10-12)**. A mega-scale introgression was identified in both the Central and Southern China at around 20-30 Mb on chromosome 6, covering a key flowering time gene *E1* (**fig. S10 and 12**). By further examining the outlier windows with the top 5% *f*_d_ values, we found introgression regions that were in common among the three geographic regions (**fig. S13**). The super exact test(*20*) revealed that the sharing of introgression regions among populations was significantly enriched (*p* < 0.0001) at any combination (**fig. S13**), indicating that variation at key common loci was important for its spread both south and north.

### Genomic signals during the spreading of soybean

Based on the phylogenetic, population structure and demographic analyses, we propose an evolutionary route of the wild and cultivated soybeans that includes four geographic paths (**Fig. 3A).** The first path corresponds to the expansion of the wild soybean from the Southern to the Northern China. The second represents the domestication process in Central China while path three is the expansion of the landrace populations from the Central region to the north and south and the fourth reflects the improvement process. We then identified signatures of selection for each of the four paths with three statistics: log_2_ (*θπ* ratio)(*21*), Population Branch Statistics (PBS)(*22, 23*), and cross-population composite likelihood ratio (XP-CLR)(*24*). Considering that the low overlap of selected molecular markers could have resulted from different signatures of population variations in the three methods(*25, 26*), we took the windows with support from at least one statistic as the candidates (**Fig. 3B and fig. S14-20**).

**Fig. 3.**
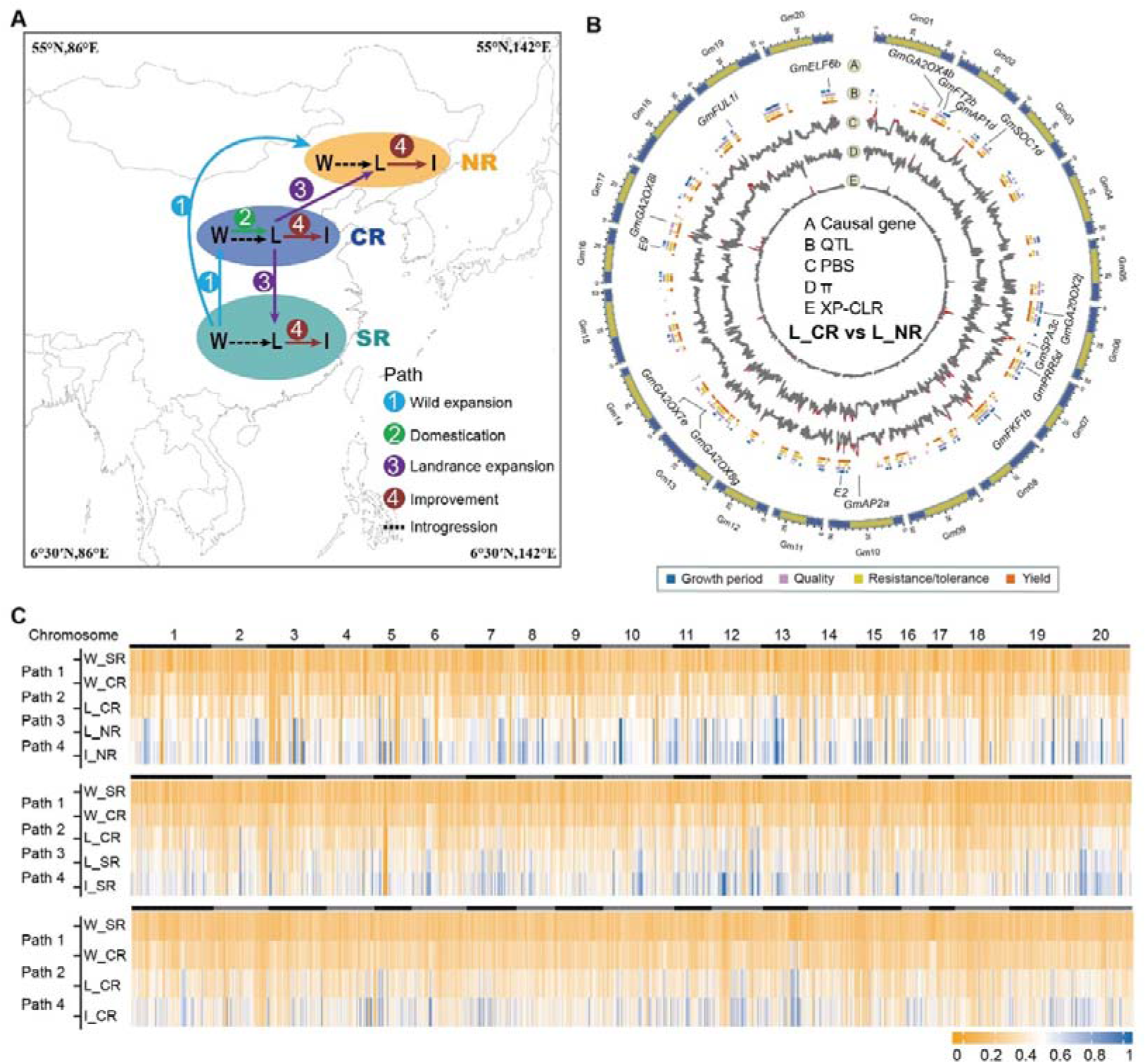
Detection of genomic regions and genes with selection signals during domestication, expansion, and improvement of soybean. (**A**) Four potential evolutionary routes of soybeans. The dashed black lines indicate the gene flow event detected by TreeMix. The numbers on the solid line represent the four routes suggested in the main text. (**B**) Genome-wide cross-population selection signatures in L_CR compared with L_NR on the basis of 8,785,134 informative SNPs. The outermost ring indicated 20 chromosomes. Functional genes with selection signals were marked in ring A, and QTLs associated with growth period, seed quality, yield and resistance/tolerance were shown in ring B with different colors. Rings C to E represented selective signals for PBS, *θπ* ratio and XP-CLR, respectively. The genomic region with selection signal (the threshold value was set as Top 1%) were indicated in red. (**C**) Heat map of allele frequency change across integrating putative intact evolutionary stages in Northern (top), Southern (mid), and Central regions (bottom). Allele frequency of each population represented population mutation frequency by using the major genotypes of the southern wild soybean population as reference. Abbreviations: “W_” represented the wild soybean, “C_” denoted the cultivated soybean, “L_” represented the landraces and “I_” denoted the improved cultivars; SR indicated the Chinese Southern region; CR implied the Chinese Central region surrounding the mid-down stream of Yellow River valley; NR stand for the Chinese Northern region plus Japan, Korean peninsula and Russian Far East region.

We identified a total of 2,438 and 4,877 genes exhibiting strong genetic differentiation during the expansion of wild soybeans (W_SR vs. W_CR, W_SR vs. W_NR) and landraces (L_CR vs. L_NR and L_CR vs. L_SR) (**Fig. 3B, fig. S14-27 and table S6**), respectively. As the wild and landrace populations have a similar geographic range, we further asked whether both populations underwent parallel adaptation to similar environments by inspecting the overlap of selected genomic regions. The genomic scan revealed that 72.94 Mb and 61.35 Mb of the genome was selected in landraces and wild relatives, respectively. Of this, 6.96 Mb was shared, which was not significantly over-represented (*p* > 0.05, hypergeometric test). Similarly, the overlap of selected genes during the expansion of the *G. soja* and *G. max* was significantly lower than expected (*p* > 0.05, hypergeometric test) by chance. Taken together, this suggests that *G. soja* and *G. max* adapted independently to the same/similar local environments. We did not find any previously characterized genes with signals of selection in path one, likely due to the limited studies in *G. soja*. In path three, a cloned flowering time gene *E2*(*27*), a homologue of GIGANTEA (GI) in *Arabidopsis*, was shown to be under selection when landraces dispersed from the Central to the Southern and Northern regions, respectively; whereas another flowering time gene *GmFT2a*, homologue of *Flowering Locus T* (*FT*) in *Arabidopsis* was detected as an outlier when the landraces expanded from the Central to the Southern region(**table S6**). Analyses of the causal variant in *E2* (Chr10:45310798) revealed that the early flowering allele (Chr10:45310798_T) was nearly fixed in the C_NR, including L_NR and I_NR. This indicates that flowering time genes are essential to the local adaptation during the geographic expansion of landraces.

We then focused on the domestication (W_CR vs. L_CR) and improvement processes (L_CR vs. I_CR; L_SR vs. I_SR; L_NR vs. I_NR), paths two and four (**fig. S16, 18-20 and table S6)**. During domestication, in order to pursue the rapid and uniform seed germination as well as a safe and edible soybean seeds without hazardous allergens(*28*), the early farmers focused on phenotypic changes that led to the loss of seed hardiness and the loss of seed bloom. In total, 2,496 genes were detected with signatures of selection during the process of domestication. Among those genes, we found several genes responsible for key phenotypic changes during domestication, such as the flowering time genes *E4* and *GmFT5a*. In addition to these candidate genes identified with a stringent cutoff (above the 99% quantile), other candidate genes appeared when the cutoff was lowered to the top 5% outliers, including a seed blooming *Boom1* (*B1*), seed hardness *GmHs1_1*, and seed dormancy *G* gene (*28-30*). The human-favored causal alleles of these genes were strongly selected in the landraces (**fig. S21**).

As breeders mostly utilized local landraces to develop improved cultivars suitable for local environments, we tested the genes under selection in three independent improvement processes. We found 2,529, 2,785, and 2,388 candidate genes in the Northern, Central, and Southern regions, respectively. Of these, 86.9% (5,921 genes) were region-specific, which may be attributed to the distinct improvement intention and the different environmental conditions in the three geographic regions. For example, three previously characterized flowering time genes exhibited selection signals during the improvement process. Among the three flowering time genes, *E1* was detected in Southern region, *E2* in Southern and Northern regions and *GmFT5a* in Central region.

To complement the three previous statistics of selection, we also identified stepwise and directional increase or decrease of allele frequencies through integrating combinations of the four paths incorporating domestication, expansion, and improvement in the three regions (**Fig. 3C**). A total of 543, 1,444 and 3,487 genes exhibited consecutive dynamic changes of allele frequencies with respect to evolutionary stages in the Southern, Northern, and Central regions, respectively. Two hundred genes were shared, suggesting their significance in the broad sense of soybean domestication. Interestingly, three previously cloned flowering time genes (*GmFT2b, GmFT4*/*E10* and *GmGBP1*) showed consecutive increases in allele frequencies. These three genes all had exhibited selection during the expansion of wild soybeans from the Southern to Central regions and domestication in the Central region (**fig. S22**). Subsequently, *GmFT2b* and *GmFT4* were utilized by the breeders during improvement in the Central regions, and *GmGBP1* and *GmFT2b* were selected during range expansion of landraces from the Central to the Northern and Southern region, respectively and then utilized in local improvements. In conclusion, those findings represent a small and stepwise, directional and consecutive shift of allele frequencies during soybean domestication, complementing the pronounced shift of allele frequencies detected by the previous statistics and providing additional genetic insights into the process of domestication and resources for further improvement.

### Validation of flowering time gene *GmSPA3c* involving in soybean expansion

As flowering time is a key agricultural trait for its contribution to adaptation, crop yield and quality, it has been the long-term target of selection during breeding research (*31, 32*). Although several flowering time genes (such as *E1-E4*, *GmFT1a*, *GmFT2a*, *GmFT2b, GmFT4, GmFT5a*, *J*, and *GmPRR3b*/*Tof12*) have been characterized in soybean (*9, 27, 33-41*), the underlying molecular mechanisms in soybean evolution remain unclear. Among the selected genes during domestication, improvement, and expansion of soybean, we observed seven cloned flowering time genes in soybean, including *E1*, *E2*, *E4*, *GmFT2a, GmFT2b, GmFT4* and *GmFT5a*.

We further found that four *FT* paralogous genes (*GmFT2a, GmFT2b, GmFT4* and *GmFT5a*) associated with photoperiod response were selected in different evolutionary paths. *GmFT5a* underwent selection during domestication and improvement, whereas *GmFT2a* was selected during landrace range expansion (**table S6**). *GmFT2b* and *GmFT4* underwent a consecutive allele frequency change from path one to four, suggesting its significance in the broad sense of domestication. Recently, the homozygous quadruple mutant with loss-of-function mutations in the four copies of MADS-box transcription factor *GmAPETALA1* genes (*AP1a-d*) was generated using CRISPR-Cas9 technology, which exhibited delayed flowering under short days in soybean (*42*). Our results indicated that *GmAP1c*, *1d* and *1b* were selected during wild range expansion, landrace expansion, and genetic improvement, respectively. To summarize, the flowering time pathway played a significant role in the broad adaptation of soybean and the paralogs of flowering time genes were selected in different evolutionary paths, suggesting the specificity of the pathway in soybean.

Of the identified candidate genes, 203 were previously described as either regulators or homologs of flowering-time genes, which covered the primary components of the flowering time pathways (**table S6**). To establish correlations between these candidate flowering genes and the genetic regions controlling flowering time, we performed a genome-wide association study (GWAS) for flowering time with 1,993 cultivated soybean lines planted in Nanjing city (32.07°N, 118.78°E). Of the 35 candidate QTL regions with association signals (−log_10_ *p* ≥ 13), a locus *qFT06-5* corresponded to the *E7* locus (**Fig. 4A, fig. S23, table S7**), one of 12 major flowering loci (*43*) and mapped in a 12.56 Mb genomic region (Chromosome 6: 31,490,622-44,050,041) flanked by two SSR markers Satt100 and Satt460 (*44*). In order to identify the candidate flowering time gene(s), we delineated a 271.8 kb region (chromosome 6: 39,983,666-40,255,433) with *r^2^* ≥ 0.8 flanking the locus *qFT06-5*. This region was under selection during the expansion of landraces from the Central to Northern regions and improvement in the Central region (**fig. S24**). Five annotated genes were located within *qFT06-5*, including two homologous genes of *Suppressor of PHYA-105* (*SPA*) (*Glyma.06G241900* and *Glyma.06G242100*). *Glyma.06G241900* was an incomplete gene due to the deficiencies of the open reading frame, while *Glyma.06G242100* is one of the four co-orthologs of *Arabidopsis SPA3,* hereafter, *GmSPA3c* (**fig. S25**).

**Fig. 4.**
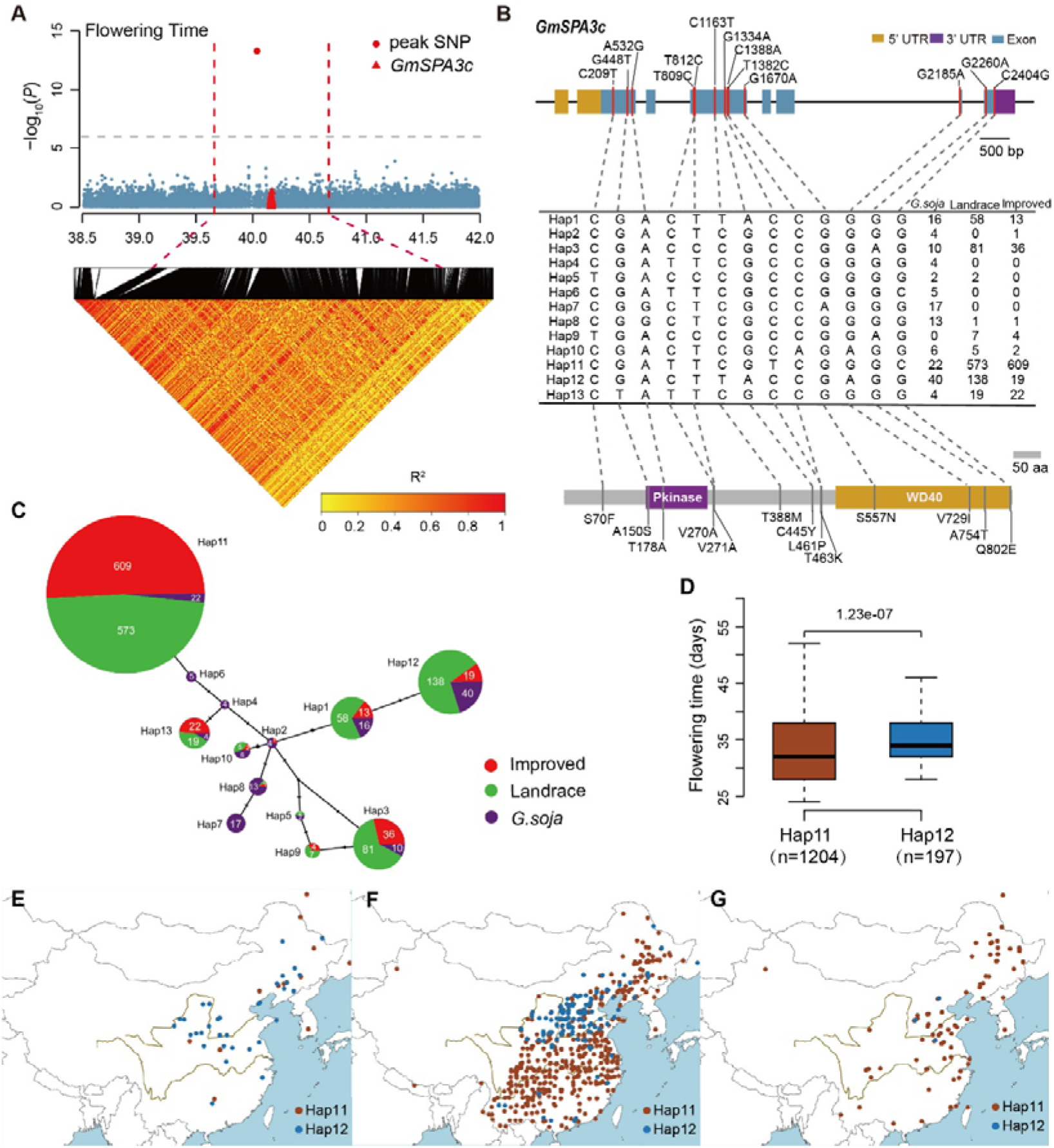
Identification of *GmSPA3c* in regulating flowering time by a genome-wide association study and characterization of the evolution and geographical distribution of *GmSPA3c* haplotypes. (**A**) Manhattan plots for the flowering time measured in the Nanjing station. Dashed red lines specified the candidate region surrounding the GWAS hit SNP. Heatmap underneath showed the LD block of SNPs in the region. Red triangles indicated the location of *GmSPA3c*. (**B**) The gene and protein models, and haplotype diversity of *GmSPA3c*. The gene model in the top panel demonstrated the UTRs (black rectangles), CDS regions (teal rectangles), and introns (horizontal solid black lines). The vertical solid lines represented the SNP loci in all the soybean samples. The protein model in the bottom panel demonstrated the structure of GmSPA3c containing the Pkinase and WD40 domains. The vertical solid lines represented the corresponding change of amino acid. (**C**) Median-joining network of thirteen *GmSPA3c* haplotypes. The pie charts in different colored were for *G. soja*, landraces, and improved cultivars, respectively. The different colored portions in each pie chart represented the number of accessions of different haplotypes. (**D**) Boxplot of the flowering time (days) of the indicated haplotype groups in the Nanjing station. The significant different levels (one-way ANOVA analysis) were showed above the bars. (**E-G**) The geographical distribution of 1,993 soybean accessions carrying different haplotypes in *G. soja* (**E**), landraces (**F**) and improved cultivars (**G**), respectively.

We detected 13 *GmSPA3c* haplotypes based on 13 missense mutations (referred to as Hap1 to Hap13). The haplotype diversity decreased from 13 in *G. soja* to 7 in landrace and 6 in improved cultivars (**Fig. 4B**). Among them, Hap11 and Hap12 are the two predominant haplotypes within cultivated soybeans including the 24 accessions with released *G. max* genomes (*2*). Median-Joining network analysis suggested that Hap11 and Hap12 were independently selected during domestication, but Hap11 rather than Hap12 was preferentially selected during genetic improvement (**Fig. 4C**). We further evaluated the phenotypic effects of the two haplotypes and revealed that the cultivars carrying Hap11 flowered significantly earlier than those carrying Hap12 (**Fig. 4D**). The geographical distribution of the accessions carrying Hap11 or Hap12 indicated that the landraces or cultivars carrying Hap11 were distributed all over China, while those carrying Hap12 were mainly distribute in central China. Together, the early flowering effect of Hap11 may contribute to a beneficial trait that was preferentially selected and expanded the adaptability of *G. max* to different regions.

We evaluated the function of *GmSPA3c* Hap11 by the CRISPR/Cas9 method (**fig. S25B**). We obtained multiple independent lines and selected two representative null mutants for phenotypic analyses. The results showed that loss-of-function of *GmSPA3c* conferred early flowering phenotype under long-day but not short-day conditions (**Figs. 5A and 5B**). Consistent with this, the flowering inhibitor *E1* and *GmFT4* were downregulated (**Fig. 5C**), while the flowering activator *GmFT5a* and *GmFT2a* was upregulated in the *Gmspa3c* mutants in a long-day photoperiod dependent manner (**Fig. 5C**), but their expressions did not change obviously under short-day conditions (**fig. S25C**). The observation that the *Gmspa3c* mutants flower earlier than the wild type TL1 (carrying Hap11) suggested that Hap11 is still a flowering repressor but with a weaker activity in comparison to Hap12.

**Fig. 5.**
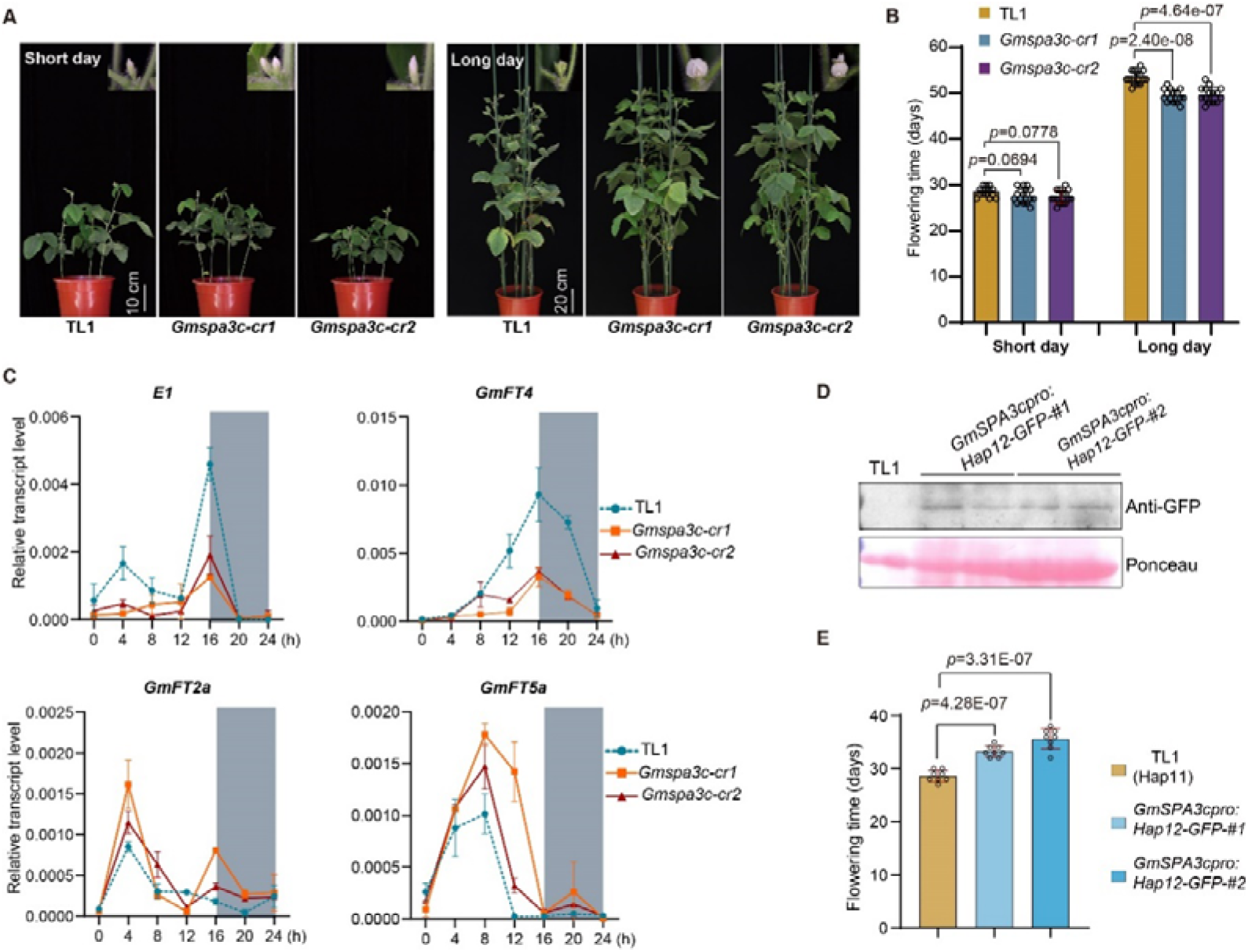
Confirmation of *GmSPA3c* as a flowering repressor and comparing the activities of Hap11 and Hap12 to regulate flowering time. (**A**) The representative images of the *Gmspa3c* mutants and wild type TL1 grown under short day (12 h Light / 12 h Dark) and long day (16 h Light / 8 h Dark) conditions for 28 and 50 days post seed germination, respectively. (**B**) Bar plot showing the flowering time of the *Gmspa3c* mutants and TL1 as in (a). Mean values ± s.d. are shown. The significant differences were determined by Student’s t-tests (n > 10). (**C**) The dynamic transcriptional level of *E1, GmFT4, GmFT2a, GmFT5a* in the *Gmspa3c* mutants and TL1 under long day conditions. Mean values ± s.d. (n = 3) are shown. (**D**) Immunoblots show the abundance of GmSPA3c-Hap12-GFP fusion proteins in the *GmSPA3cpro*:*GmSPA3c*-Hap12-GFP transgenic plants using anti-GFP antibody. The TL1 sample was used as the negative control. The proteins recognized by ponceau were used as the loading control. (**E**) Comparison of the flowering time between TL1 and the *GmSPA3cpro:Hap12-GFP* transgenic lines (Student’s t-tests; n=8). Two independent transgenic lines were generated by genetic transformation of the *GmSPA3c-Hap12* CDS driven by its 2064 bp native promoter into TL1.

To gain insight into how Hap12 is functionally different from Hap11, firstly, we compared the expression levels of *GmFT2a* and *GmFT5a* in accessions harboring either Hap11 or Hap12 (**fig. S25D**). The results showed that the expression levels of *GmFT2a* and *GmFT5a* were significantly higher in the Hap11 accessions than in the Hap12 accessions, indicating that Hap11 might be selected as a weaker flower inhibitor during soybean domestication and improvement. Besides, we performed correlation analysis between *GmSPA3c* (Hap11 and Hap12) and *GmFT5a* mRNA levels in calluses transformed with *35s:GmSPA3c* (Hap11 and Hap12) at ZT4. The results showed that expression levels of *GmFT2a* and *GmFT5a* were more effectively suppressed by Hap12 than Hap11(**fig. S25E**). Furthermore, we performed complementation experiment by genetic transformation of the *GmSPA3c*-*Hap12* CDS driven by its native promoter into TL1 (Hap11 background) (**Figs. 5D**), The results showed that the flowering time of complementation lines were significantly later than TL1, even under short-day conditions **(Fig. 5E).** The flowering inhibitor *E1* were upregulated, while the flowering activator *GmFT5a* and *GmFT2a* was downregulated in the complementation lines (**fig. S25F**).Those results that *GmSPA3c* is a flowering repressor and the behavior of its loss-of-function mutant is reminiscent to that of the *e7* NIL lines (*45*), together with the fact that the *GmSPA3c* gene locates within the *E7* QTL site, suggested that *GmSPA3c* is a *bona fide* candidate for the long sought-after *E7* gene.

## Discussion

Distinct from previous publications which majored on the studies of soybean domestication and genetic improvement related to artificial selection (*2, 4-6*), this study provides the first comprehensive analyses of the evolutionary history of soybean, including the dispersal of wild soybean, domestication site, range expansion of landraces and the subsequent improvement process, based on a dense and diverse sampling of the wild and cultivated samples. *G. soja,* the wild progenitor of cultivated soybean, has not undergone artificial bottlenecks and thus, is one valuable genetic source with ability conferring adaption to new environments (*19*). To improve the study and utilization of *G. soja*, we constructed the phylogenetic tree using the only two perennial species (*G. tabacina* and *G. tomentella*) discovered in East Asia (*46*) as root and deduced the phylogeographical expansion routes of wild soybeans, which maybe origin in south China and spread to central and north China. Suggested that even though its wild progenitor originated in the subtropical Asia, soybean was domesticated in the temperate region in China. We also provide genetic evidence to support that soybean was domesticated in Central China surrounding the middle and lower reaches of Yellow River (*16, 47*) from its wild progenitor, which originally spread from Southern China. Post domestication, landraces expanded northward and southward colonizing an expansive area of East Asia. Recurrent introgression from adapted sympatric wild soybeans might facilitate the local adaptation of landraces. The improved soybean was most likely developed from locally adapted landraces, indicating that soybean cultivars have recurrently ‘used’ local wild genetic diversity. Given the independent adaptation between the wild and cultivated soybean, the introduction of genetic resource of wild soybean from allopatric regions would likely be valuable to mitigate its adaptation to the changing climate.

Most previous studies of crops described selection signals during domestication and genetic improvement based on the broad sense of genomic changes (*4, 48*). Here we included geography to further refine the genetic footprint of breeding within specific regions. As soybean was constrained and adapted to a narrow and specific geographic range, we detected the selection signals during domestication between the wild and landrace populations from the domestication site, and during improvement between sympatric landrace and improved lines, which minimize noise resulting from the complex genetic background of the contrast population. Additionally, the traditional methods are all based on the comparison of two different populations (*21, 23, 24, 49*). However, the domestication, expansion and improvement are consecutive evolutionary processes. So, we designed a method to detect stepwise and directional increase or decrease of allele frequencies across multiple evolutionary stages (*50, 51*), which provides insights into the evolutionary dynamics of soybean genomes under the combination of both natural and artificial selection. Given the importance of flowering time in the spread and adaptation of soybean, we evaluated candidate genes in the flowering time pathway. We used an integrated strategy by combining the search for genomic footprints of selection with association mapping to identify suitable candidate genes. The most important and laborious step was the validation of candidate genes by generating loss-of-function mutants by CRISPR/Cas9 technology and complementation experiment. This strategy is in particular suitable for functional variant encoded by single nucleotide polymorphisms. Extending our approach towards structural variation is challenging, but as a first important step an integrated graph-based genome for soybean using *de novo* assembled genomes of 29 genotypes has been published(*2*).

Our results highlight that the adaptation of flowering time was a continuous process. In particular, we verified the flowering time modulating functions of one selected gene, *GmSPA3c* as a flowering repressor, underwent weak but sustained selection during domestication, landrace expansion and improvement. Our GWAS mapping and functional verification further suggested that *GmSPA3c* is a candidate of long-sought flowering locus *E7*. *GmSPA3c* featured early flowering alleles or haplotypes of both genes were selected, which is consistent with a historic trend that novel varieties flowered earlier locally since the beginning of domestication (*52*). The shortening of the time from vegetative to reproductive growth may associate with the climate change or be preferred by the breeders to ensure the harvestability of seeds (*53*). However, the intervals between flowering and maturity tended to become longer which may contribute to the production enhancement(*54*). In summary, our study not only shed lights into the evolutionary history of soybean, but also provides valuable genetic resources for future breeding.

## Materials and Methods

### Plant materials and growth conditions

A total of 2,214 soybean accessions including cultivated *G. max* (1,993), annual wild *G. soja* (218), perennial wild species *G. tomentella* (2) and *G. tabacine* (1), were analyzed in this study (table S1). Among them, 1,674 genomes were newly sequenced in this study and the rest 540 have been released before(*55, 56*). *G. tomentella* and *G. tabacine*, the only two perennial wild species occurring in China, were included as out-groups for the population structure analyses. Most (99.5% in 218 accessions) of *G. soja* were selected from its native range (Ease Asia), including China (179), Korea (10), Japan (19) and Russia (9), to well represent the diversity of this species. Among 1993 *G. max* accessions, 1,131 were landraces mainly selected from Chinese primary and applied core collections to capture as much diversity of the 23,587 cultivated soybean accessions preserved in the Chinese National Soybean GeneBank as possible. The rest 862 improved cultivars were collected from 17 countries, mainly from the main soybean producing countries such as United States, China, Japan and Korea (table S1). Of 218 *G. soja* accessions, 109 were planted in two experimental fields, Jingzhou city in Hubei province (30.3 °N, 112.2 °E) in 2014, and Beijing city (40.1 °N, 116.7 °E) in 2015. Experiments were performed using a completely randomized experimental design with two complete replicates in Jingzhou and three complete replicates in Beijing. Moreover, 1,498 *G. max* accessions were planted in Nanjing city, Jiangsu province (32.0 °N, 118.8 °E) in 2018. Flowering time was scored based on the description in Qiu et al. (2006)(*57*).

### DNA isolation and genome sequencing

The genomic DNA was extracted with a total amount of 1.5 μg per sample and used as input material for the DNA sample preparations. Sequencing libraries were generated using TruseqNano^®^ DNA HT sample preparation Kit (Illumina USA) following manufacturer’s recommendations and index codes were added to attribute sequences to each sample. Basically, the libraries were prepared following these steps: the genomic DNA sample was fragmented by sonication to a size of ~350 bp, then DNA fragments were end-polished, A-tailed, and ligated with the full-length adapters for Illumina sequencing with further PCR amplification. At last, PCR products were purified (AMPure XP bead system) and libraries were analyzed for size distribution by Agilent2100 Bioanalyzer and quantified using real-time PCR. Subsequently, we used the Illumina Hiseq X platform to generate ~10.58 Tb raw sequences with 150-bp read length. Additionally, 540 were previously released accessions with 5.94 Tb sequences were download from NCBI database and incorporated to analysis.

### Sequence quality checking and filtering

To avoid reads with artificial bias, i.e. low-quality paired reads, which primarily result from base-calling duplicates and adaptor contamination, we removed the following types of reads: (i) reads with ≥10% unidentified nucleotides (N); (ii) reads with >10 nt aligned to the adaptor, with ≤10% mismatches allowed; (iii) reads with >50% bases having phred quality <5; and (iv) putative PCR duplicates generated through PCR amplification in the library construction process, i.e. read 1 and read 2 of two paired-end reads that were completely identical. Consequently, we obtained 16.41 Tb (~ 6.3X coverage per individual) of high-quality paired-end reads, including 96.05% and 90.98% nucleotides with phred quality ≥ Q20 (with an accuracy of 99.0%) and ≥ Q30 (with an accuracy of 99.9%), respectively (table S1).

### Sequence alignment, variation calling, and annotation

After sequence quality filtering, we first mapped the remaining high-quality sequences to the soybean reference genome(*12*) (Williams 82 assembly V2.0, http://www.phytozome.net/soybean) using BWA software (v. 0.7.17-r1188)(*58*) with the command ‘mem −t 10 −k 32 −M’. Second, we converted SAM format to BAM format using the package SAMtools (v.1.3)(*59*). Third, we sorted BAM files using the package Sambamba (v. 0.6.8)(*60*). Finally, the sorted bam file was marked as duplicate using the command “MarkDuplicates” in the package picard (v. 2.18.15, http://broadinstitute.github.io/picard). Subsequently, we performed individual gVCF calling according to the best practices using the Genome Analysis Toolkit (GATK, version v4.1.2.0)(*61*) with the HaplotypeCaller-based method and then population SNP calling by merging all gVCFs with the commands “GenomicsDBImport” and “GenotypeGVCFs”. Consequently, a total of 65,374,688 SNPs (60,153,828 are bi-allelic) and 10,952,749 indels (8,349,613 small insertions and deletions < 15 bp and less than 50% missing) were identified in 2,214 accessions.

To obtain credible population SNP sets, we performed a screening process as follows:

a. For filtering SNPs, the hard filter command ‘VariantFiltration’ was applied to exclude potential false-positive variant calls with the parameter ‘--filterExpression “QD < 2.0 ∥ MQ < 40.0 ∥ FS > 60.0 ∥ SOR > 3.0 ∥ MQRankSum < −12.5 ∥ ReadPosRankSum < −8.0”’. Subsequent filtering was performed after removing three perennial wild accessions.
b. Screening of biallelic variants was performed with a Hardy–Weinberg equilibrium p-value >= 0.01(*62*).
c. Variants were filtered out when the proportion of samples within the population lacking the variant was > 20% and the minor allele frequency (MAF) was < 0.01.
d. Variants were filtered out when the inbreeding coefficient was more than 0.348(*63*). After those steps, we obtained 8,785,134 high-credible biallelic SNPs,
e. Subsequently, we also subsampled 1,721,062 SNPs set using a two-step linkage disequilibrium (LD) pruning procedure with PLINK (v1.9) in which SNPs were removed with a window size of 10 kb, window step of one SNP and r^2^ threshold of 0.8, followed by another round of LD pruning with a window size of 50 SNPs, window step of one SNP and r^2^ threshold of 0.8. Thus, these 1,721,062 SNPs is used for subsequent population structure analyses in soybeans.
f. We added genotypes of three perennial accessions to the filtered SNP set of 2211 accessions. The integrated SNP data set was used to infer population structure of 2214 accessions.

### Annotation of genomic variants

Genomic variant annotation was performed according to the soybean genome using the package ANNOVAR (version: 2019-10-24)(*64*). Based on the genome annotation, genomic variants were categorized as being in exonic regions, UTR regions (represent the 5’ and 3’ untranslated sequences), intronic regions, splice sites (within 2 bp of a splicing junction), upstream and downstream regions (within a 2-kb region upstream or downstream from the transcription start site), and intergenic regions. The functional consequences of the variants in coding regions were further grouped into synonymous, missense, stop-gain, stop-loss.

### Population diversity statistics

We first screened out windows with more than 10 SNPs. Subsequently, nucleotide diversity (*θπ*)(*65*) was applied to estimate the degree of variability within each group and genetic differentiation (*F*_ST_)(*66*) were applied to explain population differentiation on the basis of the variance of allele frequencies between two different groups by VCFtools (v0.1.14)(*67*).

### Linkage disequilibrium (LD) analysis

To estimate and compare the pattern of LD for different groups, the squared correlation coefficient (*r^2^*) between pairwise SNPs was computed using the software PLINK (v1.9)(*68*). Regarding the LD for overall genome, the *r^2^* value was calculated for individual chromosomes using SNPs from the corresponding chromosome with parameter ‘–ld-window-r2 0 –ld-window 99999 –ld-window-kb 1000’, and then the pairwise *r^2^* values were averaged across the whole genome.

### Population structure analysis

To investigate the genetic relationships between 2214 soybeans, we constructed a phylogenetic tree using the neighbor-joining (NJ) tree with 100 bootstrap iterations based on the 1,721,062 SNPs using the TreeBest program (version: 1.92) (https://github.com/Ensembl/treebest). The population genetic structure was examined via an expectation maximization algorithm, as implemented in the program ADMIXTURE (v1.23)(*69*), through the preset the number of assumed genetic clusters K with 10,000 iterations for each run. We also conducted PCA to evaluate genetic structure using GCTA software (v1.24.2)(*70*). We also build the maximum likelihood tree for soybean populations based on 1,721,062 SNPs set using Perennial population as outgroup applying software TreeMix(*18*).

### Genome-wide selective sweep scanning

Based on 8,785,134 high-credible biallelic SNPs, several statistical methods were employed to identify genome-wide selection signals. Firstly, by comparing the pairwise *F_ST_*(*66*) between designed compared patterns with a sliding window (10-kb windows sliding in 5-kb steps), we employed Population Branch Statistic (PBS) approach(*71*) to detect incomplete selective sweeps over short divergence times. Our approach designed to take advantage of outgroup and used to identify selection targeted on the tested lineage. The PBS was calculated as follows:

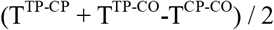

Where T represents the population divergence time in units scaled by the population size, which is the negative log transformed (1 - *F_ST_*) between two populations. TP represents the targeted population; CP indicates the control population; and CO implies the outgroup. We considered the window as the candidate selected regions when PBS value of the comparative sliding windows at a significance of *P* < 0.01 (*Z-test*).

Specifically, for comparative pattern of W_CR vs L_CR in Fig. 3b, the design formulas is (T ^L_CR-W_CR^ + T ^L_CR-W_SR^ −T ^W_CR-W_SR^) / 2; for the pattern of W_SR vs W_CR in Supplementary Fig. 25, the formula is (T ^W_CR-W_SR^ + T ^W_CR-W_NR^ −T ^W_SR-W_NR^) / 2; for the pattern of W_SR vs W_NR in Supplementary Fig. 26, the formula is (T ^W_NR-W_SR^ + T ^W_NR-W_CR^ −T ^W_SR-W_CR^) / 2”; for the pattern of L_CR vs L_NR in Supplementary Fig. 27, the formula is (T ^L_NR-L_CR^ + T ^L_NR-W_CR^ −T ^L_CR-W_CR^) / 2; for the pattern of L_CR vs L_SR in Supplementary Fig. 28, the formula is (T ^L_SR-L_CR^ + T ^L_SR-W_CR^ −T ^L_CR-W_CR^) / 2; for the pattern of L_CR vs I_CR in Supplementary Fig. 29, the formula is (T ^I_CR-L_CR^ + T ^I_CR-W_CR^ −T ^L_CR-W_CR^) / 2; for the pattern of L_SR vs I_SR in Supplementary Fig. 30, the formula is (T ^I_SR-L_SR^ + T ^I_SR-W_SR^ −T ^L_SR-W_SR^) / 2; for the pattern of L_NR vs I_NR in Supplementary Fig. 31, the formula is (T I_NR-L_NR + T I_NR-W_NR −T L_NR-W_NR) / 2.

Second, *θπ*(*65*) were calculated based on a sliding window (10-kb windows sliding in 5-kb steps) in two populations, A and B. The statistic log2 (*θπ*_A_ / *θπ*_B_) was then calculated with respect to A and B populations. An unusually negative value (1% outliers) suggests selection in population A, and the top positive value (1% outliers) indicates selection in population B.

Third, the test of cross-population composite likelihood ratio (XP-CLR; https://github.com/hardingnj/xpclr)(*72*) was performed with the following parameters: sliding window size, 0.01 cM; grid size, 10 k; maximum number of SNPs within a window, 100; and correlation value for two SNPs weighted with a cutoff of 0.95. The genetic distance was calculated based on a published genetic map(*73*). The windows with top 1% XP-CLR score were taken as outliers.

The genetic diversity π is a classic statistic to detect signals of selection (especially hard sweeps) by assuming that selected regions showed a reduced genetic diversity. PBS method can be viewed as a model-based extension of *F_ST_*, which was very powerful in detecting incomplete selective sweeps over short divergence times(*22*); while the XP-CLR method is able to detect ancient selective events(*24*). Thus, the three methods are compatible and complementary. As the three methods are good at detecting different selective events, we observed a low overlap of selected molecular markers(*25, 26, 74, 75*). Therefore, we took the candidates with support from at least one statistic. Subsequently, all candidate regions were assigned to corresponding SNPs and genes.

### Population directional mutation analysis

We examined the directional increase or decrease of allele frequencies though integrating putative intact evolutionary routes of soybeans incorporated with expansion, domestication and improvement in three regions. First, we identified the major genotypes of the southern wild soybean population and used it as a reference genotype. Then, the allele frequencies for SNPs were calculated for each population and then we calculated average allele frequencies within a sliding window (10-kb windows sliding in 5-kb steps). Subsequently, we characterized the changed trend of windows allele frequencies in different evolutionary stages. Finally, we screened out candidate windows which exhibited the consecutive dynamic change of allele frequencies.

### Demographic history analyses

We inferred the fluctuation of the effective population size for three inferred sub-populations of cultivated soybeans (C_CR, C_NR and C_SR) with SMC++ (v1.15.2)(*76*) based with a constant generation time of 1 years and the per-generation mutation rate as 6.1×10-9 (*77*). In order to test the introgression between the wild and cultivated soybeans, the *f*_d_ statistic(*78*) was computed based on a tree form (((P1, P2), P3), O), where P1 was fixed as the American cultivated lines and the three perennial wild species as the outgroup (O). P2 was set to each of the three geographical populations in landraces and P3 was defined to each of the three geographical populations in wild soybeans. The *f*_d_ statistic was computed in 10-kb non-overlapping windows with the python script ABBABABAwindows.py (https://github.com/simonhmartin/genomics_general). The windows with the 95% top *f*_d_ values were regarded as outliers.

### Genome-wide association study

Association tests were performed with a multi loci model, FarmCPU (v 1.02)(*79*), which iteratively utilized fixed effect model and random effect model. The top three columns of principal components, phenotypes and pseudo QTNs (Quantitative Trait Nucleotides) were added as covariates in the fixed effect model for association tests and the model can be written as:

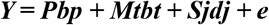

where ***Y*** is phenotypic observation vector; ***P*** is a matrix of fixed effects, including the top three principal components of all phenotypes; ***M_t_*** is the genotype matrix of ***t*** pseudo QTNs that used as fixed effects; ***bp*** and ***bt*** are the relevant design matrices for ***P*** and **M_t_**, respectively; ***Sj*** is the ith marker to be tested and ***dj*** is the corresponding effect; ***e*** is the residual effect vector and ***e ~ N(0,Iσ2e)***. Random effect model is used for selecting the most appropriate pseudo QTNs. The model is written as:

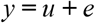

where *y* and *e* stay the same as in fixed effect model; *u* is the genetic effect and *u* ~ *N* (***0***, *K*σ***2****u*), in which *K* is the relationship matrix that defined by pseudo QTNs. In order to detect the significant SNPs, we used the Bonferroni correction threshold for multiple tests, defined as α/K (α = 0.05 and K is the number of SNPs).

### Functional verification of candidate genes

To generate the CRISPR/Cas9-engineered mutants, gRNAs were designed using CRISPR direct website (*http://crispr.dbcls.jp/*)(*80*). Multiple target gRNAs were selected for each gene to construct the CRISPR/Cas9 vector according to the protocol reported previously(*55*). The editing efficiency of each construct was evaluated by soybean hairy root system(*81*), and at least two vectors with high editing efficiency for each gene were selected for soybean transformation. The above-mentioned CRISPR/Cas9 vectors were individually introduced into Agrobacterium tumefaciens strain EHA105 via electroporation and then transformed into an elite soybean cultivar Tianlong 1 (TL1) using the cotyledon-node method(*82*). For phenotypic analysis, plants were grown under long days conditions (16 h light/ 8 h dark, 26°C), and short days conditions (12 h light/ 12 h dark, 26°C) in phytotrons. The *GmSPA3c-Hap12* CDS and the 2064 bp native promoter were obtained from Williams82 by PCR amplification. The 2064 bp native promoter and CDS fragment were amplified by overlapping PCR to obtain one fragment and then introduced into the pTF101-GFP vector replacing the *d35S* promoter-gene fragment. This new complementary construct (*GmSPA3c-Hap12*-GFP) was introduced into Agrobacterium strain EHA105, and Agrobacterium-mediated transformation of the TL1 carrying Hap11 of *GmSPA3c* was performed as described previously(*55*). All the primers used for vector construction are listed in Supplementary table 8.

### Gene expression analysis

To compare the dynamic transcriptional levels of indicated genes in the wild type, *Gmspa3c* mutant lines and complementary lines, the soybean plants were grown under long days or short days conditions for 20 days. The second fully expanded trifoliolate leaves were harvested in 4 h intervals during a 24 h period. Total RNA was extracted using TRIzol Reagent (TIANGEN) and cDNA was synthesized from DNase-treated total RNA (3 μg, reaction total volume 20 ul) using a reverse transcription kit (TransGen Biotech). qRT-PCR was performed in 384-well optical plates using a SYBR Green RT-PCR kit (Vazyme) with an ABI Q7 equipment. All primers used for indicated genes were listed in Supplementary table 8. Three independent biological replicates were performed, and three replicate reactions were employed for each sample.

### RICE System to Investigate Gene Expression

The *d35s* droved *GmSPA3c* (Hap11 and Hap12) plasmids were introduced into A. tumefaciens strain K599, which was used to infect young seedlings of Tianlong1 at the hypocotyl region to induce transgenic hairy roots according to apreviously reported method (*83*). RICE System performed as described previously(*55*). The transgenic roots were grown on the callus induction medium for 2 weeks under long-day conditions. Those independent transgenic callus lines confirmed by qRT–PCR were transferred to fresh callus induction medium for subculturing. Correlation analysis between *GmSPA3c* (Hap11 and Hap12) and *GmFT2a/5a* mRNA levels in calluses transformed with 35s droved *GmSPA3c* (Hap11 and Hap12) at ZT4 in long-day conditions.

### Accession numbers

Gene Sequences were downloaded from the *Glycine max* Wm82.a2.v1 (Soybean) database (https://phytozome.jgi.doe.gov/pz/portal.html). The accession numbers are *GmSPA3c* (*Glyma.06G242100*), *GmFT2a* (*Glyma.16G150700*), *GmFT5a* (*Glyma.16G044100*), *E1* (*Glyma.06G207800*), *GmFT4* (*Glyma.08G363100*) and *GmActin* (*Glyma.18G290800*).

## Acknowledgments

This work was partially supported by the National Key R&D Program of China(2021YFD1201601, 2016YFD0100201, and 2020YFE0202300), the National Natural Science Foundation of China (32072091), the Platform of National Crop Germplasm Resources of China (2016-004, 2017-004, 2018-004, 2019-04 and 2020-05), Crop Germplasm Resources Protection (2016NWB036-05, 2017NWB036-05, 2018NWB036-05 and 2019NWB036-05), the Agricultural Science and Technology Innovation Program (ASTIP) of Chinese Academy of Agricultural Sciences (CAAS-ZDRW202109). We thank the Core Facility Platform, Institute of Crop Sciences, Chinese Academy of Agriculture Sciences (CAAS), for assistance with sequencing.

## Author Contributions

Y-H.L., B.L., S.T., and L.Q. conceived the study. Y-H.L., L.W., Y.G, J.R., S.A.J., B.L., S.T., and L.Q. jointly wrote the paper. H.H., Y.T., Y.G.., Z.L., R.G., Z.Y., L.Z., T.L.G. provided seeds and DNAs. Y-F.L., G.X., J.W., B.F., X.W., H.Q., W.Z., X.Y.L., D.H., R.C. collected the phenotype data. X.G., X.J. performed sequencing/SNP calling. Y-H.L., L.W., C.J., D.L., Y.H. X.K.L. performed comparative/population/evolutionary/biology analyses. C.Q., H.L., T.Z., Y.G., J.L. performed the experiments.

## Competing interests

The authors declare no competing interests.

## Data deposition and accession numbers

All whole genome sequencing data in this study have been deposited in the NCBI Sequence Read Archive under accession number PRJNA681974.

## Notes

### Competing Interest Statement

The authors have declared no competing interest.

