## Supplemental Figures S1-25 for "Genome-wide signatures of geographic expansion and breeding process in soybean"

**Supplementary Figures**

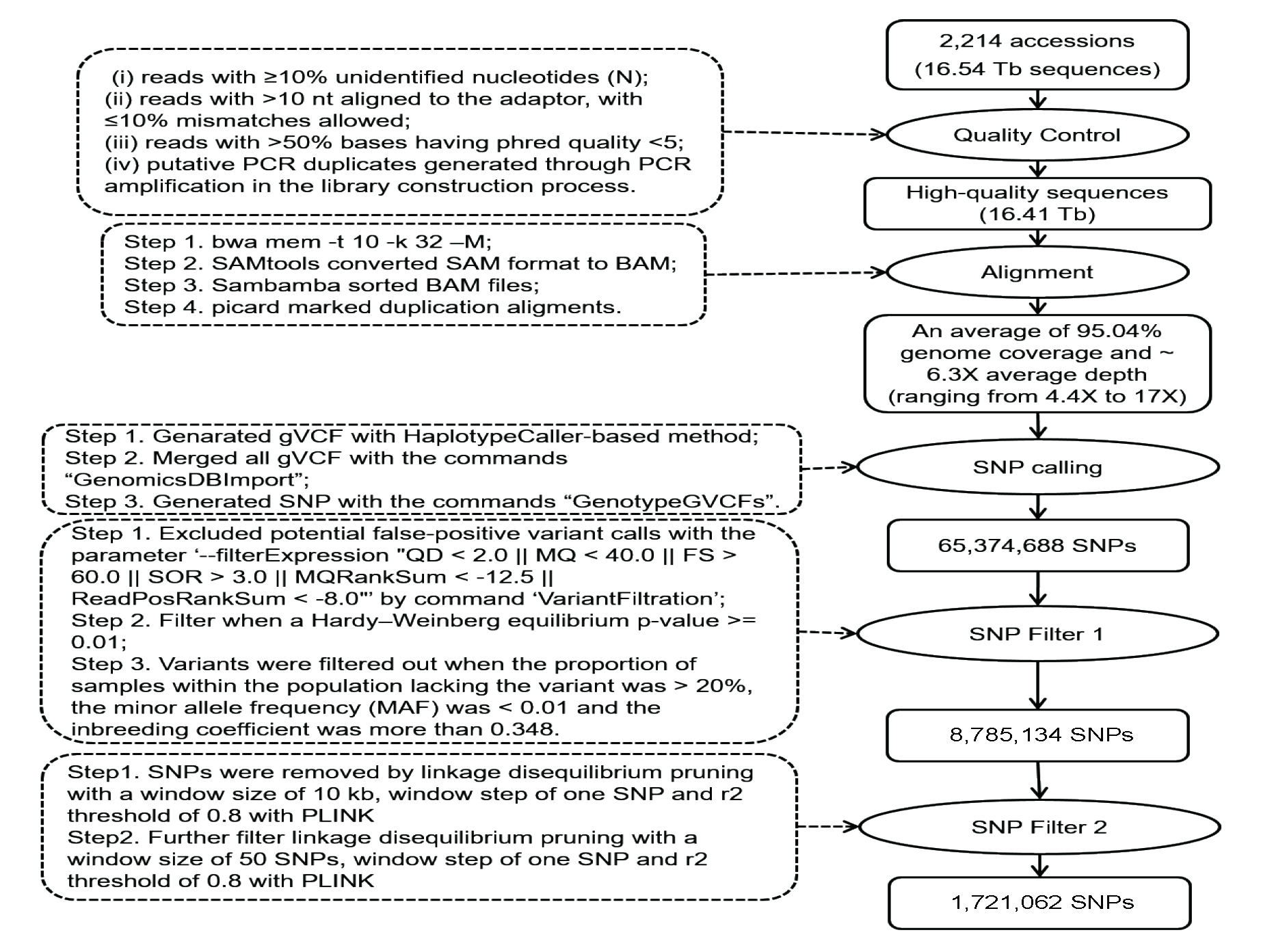

**Fig. S1. The pipeline of SNP calling and filtering in 2,214 soybean accessions**.

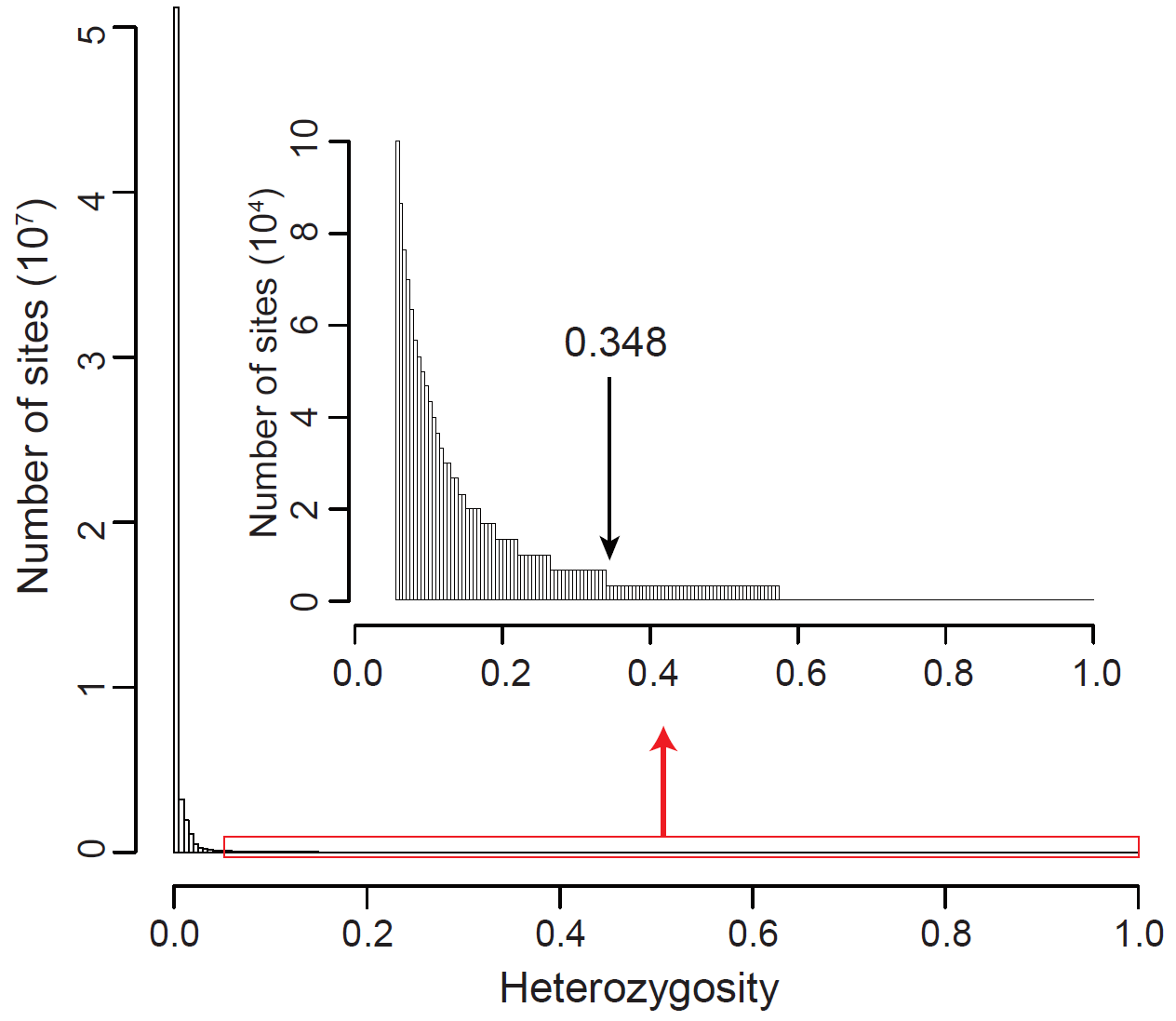

**Fig. S2. Distribution of heterozygosity of all biallelic SNPs.**

The arrow indicated the cutoff value (0.348) to exclude SNPs exceeding the expected heterozygosity based on the formula reported in Wang et al. (2018).

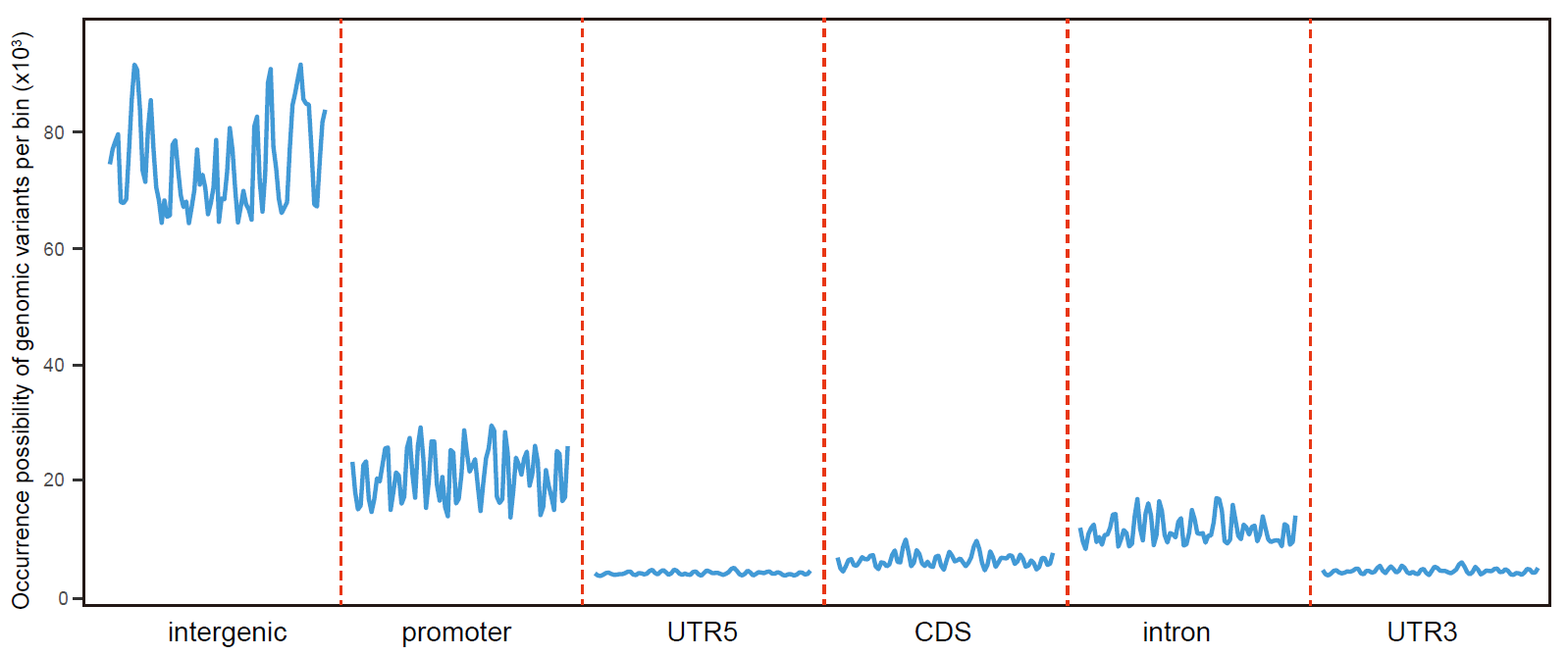

**Fig. S3. Occurrence possibility of SNPs in different genomic elements.**

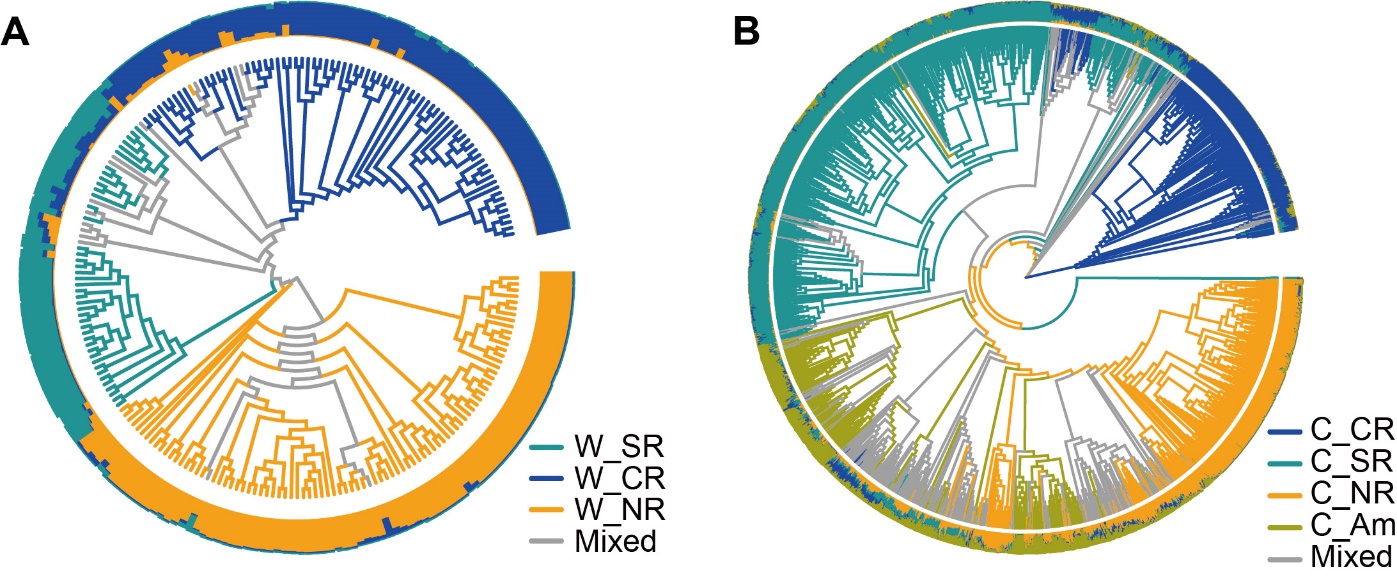

**Fig. S4. Population structure of wild and cultivated soybean accessions based on 1,721,062 SNPs.** (**A**) Population structure of 218 *G. soja* accessions. The outer ring denoted the estimated proportions of an individual’s assignment at *K* = 3. (**B**) Population structure of 1,993 *G. max* accessions. The outer ring showed the estimated proportions of an individual’s assignment at *K* = 4. Abbreviations: “W_” represented the wild soybean and “C_” denoted the cultivated soybean; SR indicated the Chinese Southern region; CR implied the Chinese Central region surrounding the mid-down stream of Yellow River valley; NR stand for the Chinese Northern region plus Japan, Korean peninsula and Russian Far East region.

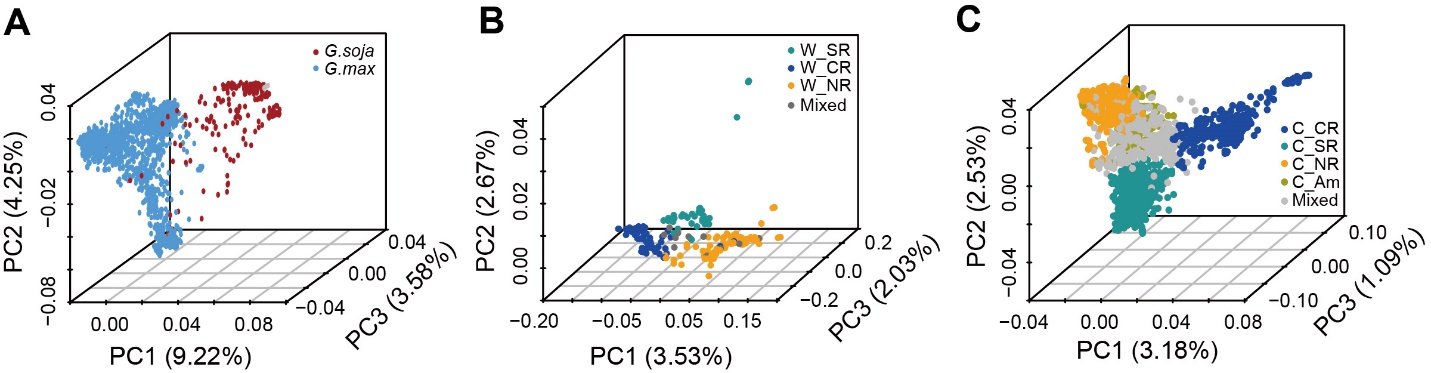

**Fig. S5. Principal Component Analysis (PCA) of 2,211 soybean accessions.** (**A**) PCA of 1,993 *G. max* and 218 *G. soja* accessions. (**B**) PCA of 218 *G. soja* accessions. (**C**) PCA of 1,993 *G. max* accessions. Abbreviations: “W_” represented the wild soybean and “C_” denoted the cultivated soybean; SR indicated the Chinese Southern region; CR implied the Chinese Central region surrounding the mid-down stream of Yellow River valley; NR stand for the Chinese Northern region plus Japan, Korean peninsula and Russian Far East region. “Am” pointed to America. “Mixed” indicated accessions with admixed genomes.

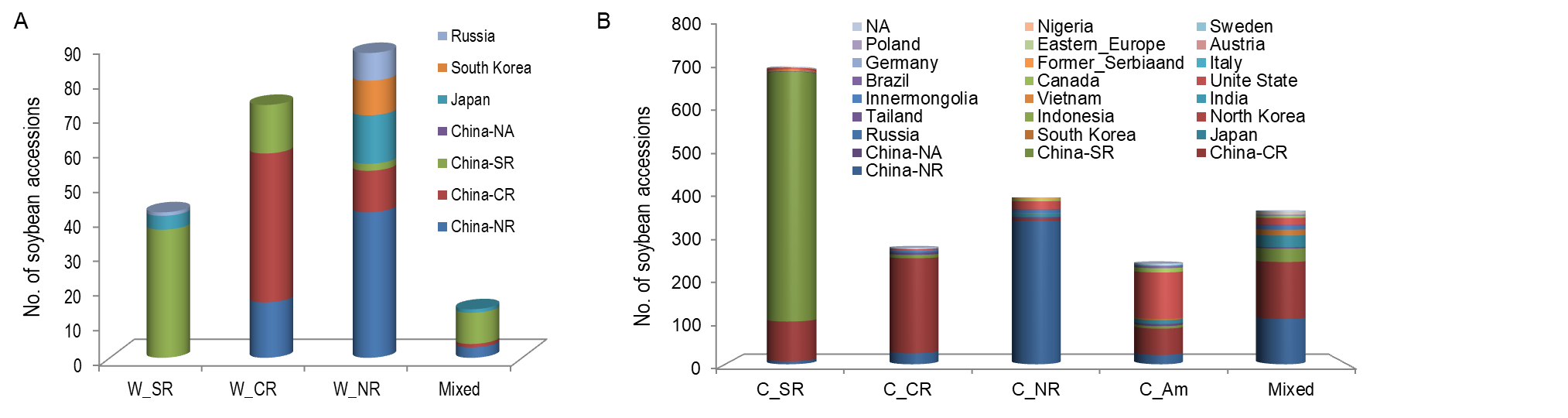

**Fig. S6. Geographic origin of the wild (A) and cultivated (B) soybeans in population structure clusters at *K =* 3 and *K =* 4, respectively.** Abbreviations: “W_” represented the wild soybean and “C_” denoted the cultivated soybean; SR indicated the Chinese Southern region; CR implied the Chinese Central region surrounding the mid-down stream of Yellow River valley; NR stand for the Chinese Northern region plus Japan, Korean peninsula and Russian Far East region. “Am” pointed to America. “Mixed” indicated accessions with admixed genomes. NA, Not avaible.

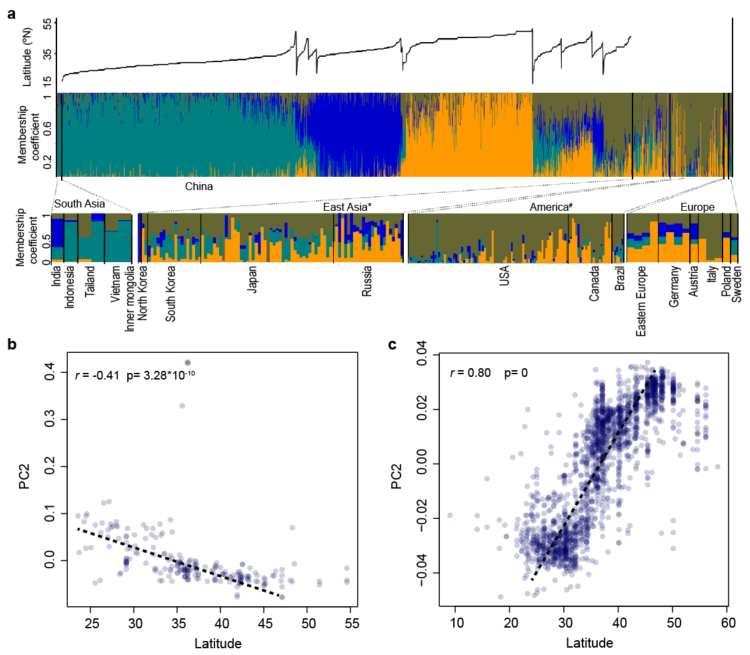

**Fig. S7. Association between population structure and latitude distribution.** Population structure of *G. max* accessions at *K* = 4 in the ADMIXTURE analysis. The accessions were ordered based on the latitude of their collection site.

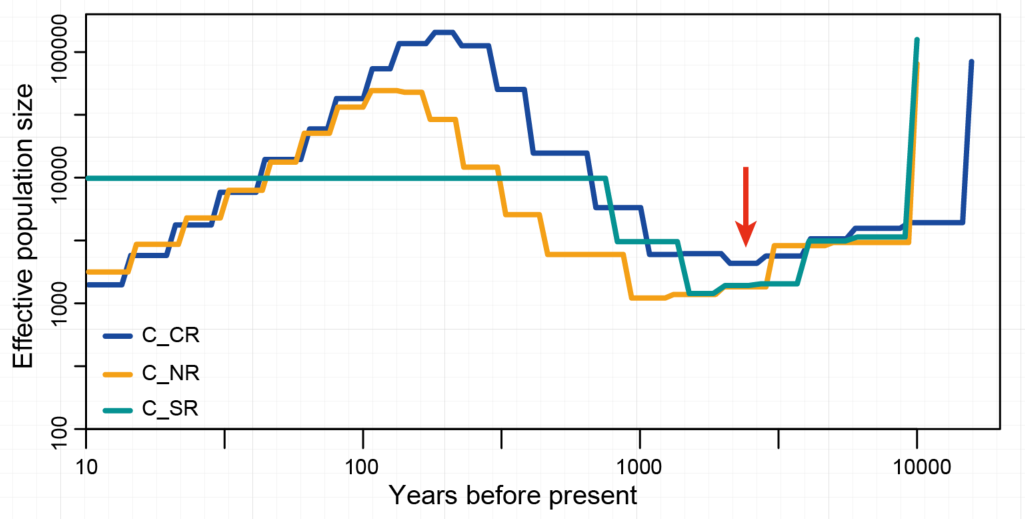

**Fig. S8. Estimates of effective population size over time in SMC++ with mutation rate of 6.1×10^-9^ and one generation per year**(*84*) **in three sub-populations of cultivated soybeans.** The x axis is log_10_ scaled. “C_” denoted the cultivated soybean; SR indicated the Chinese Southern region; CR implied the Chinese Central region surrounding the mid-down stream of Yellow River valley; NR stand for the Chinese Northern region plus Japan, Korean peninsula and Russian Far East region.

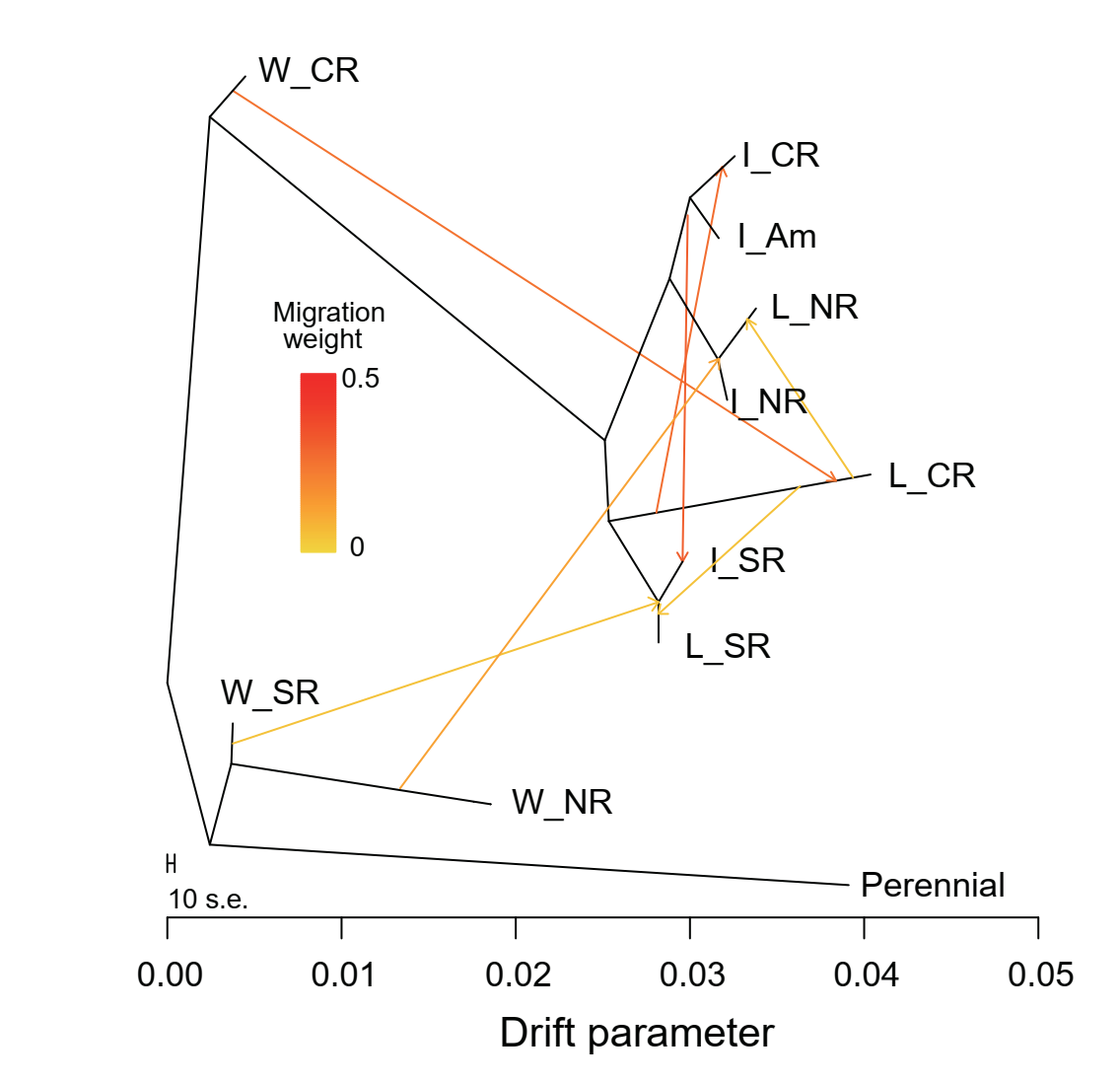

**Fig. S9.** **Population splitting and mixing analysis of 11 soybean groups in TreeMix with three perennial accessions used as the outgroup (OUT) and m=7.** The arrow indicated the migration direction. The abbreviation “W” indicates the wild soybean; “L” represents the landraces; “I” denotes the improved cultivars. SR indicated the Chinese Southern region; CR implied the Chinese Central region surrounding the mid-down stream of Yellow River valley; NR stand for the Chinese Northern region plus Japan, Korean peninsula and Russian Far East region. “Am” pointed to America.

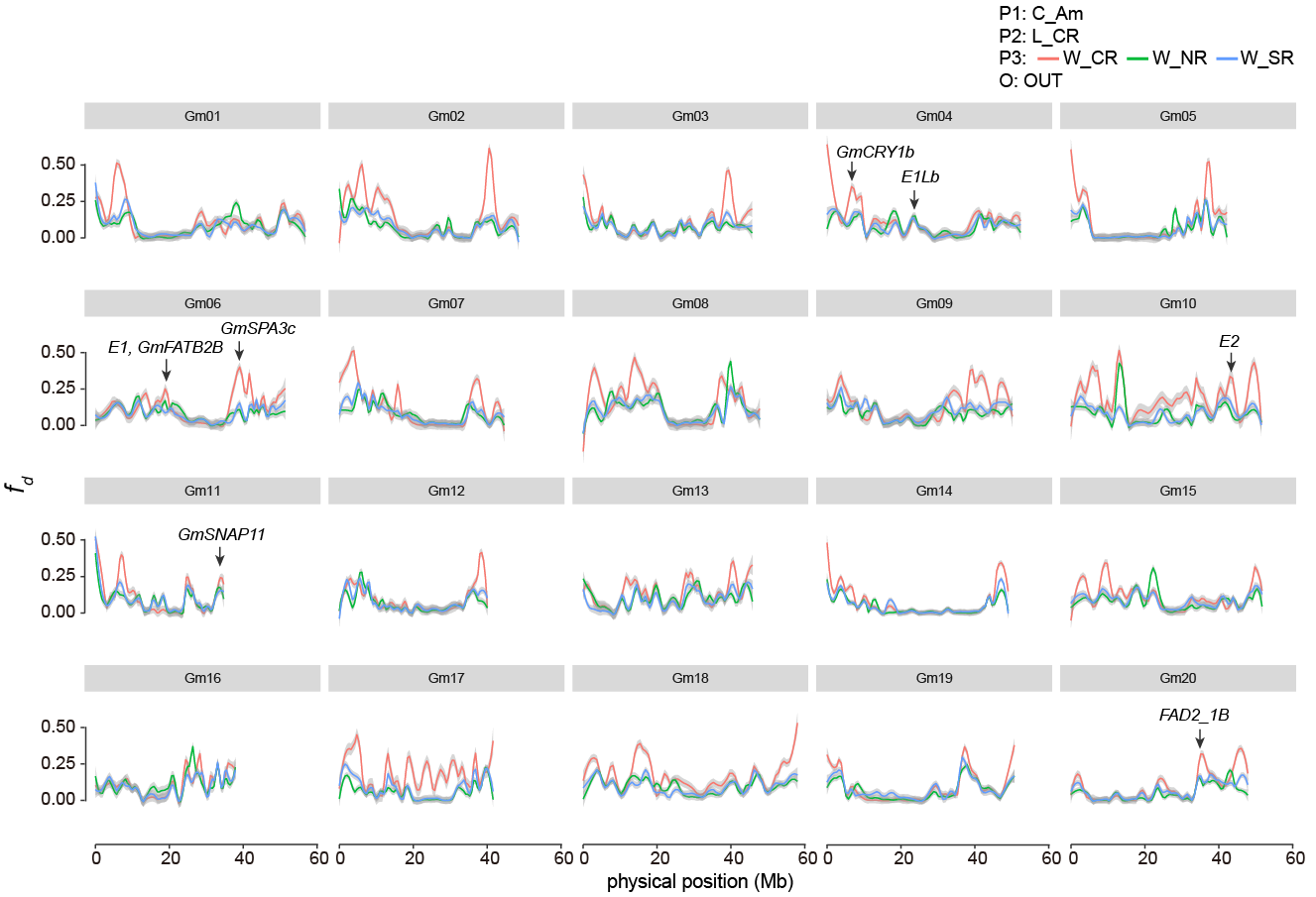

**Fig. S10.** **The distribution of *f*_d_ values in the landrace sub-population from the central region (L_CR) with potential introgression from three wild sub-populations, respectively.** The arrows denoted some characterized genes located in the outlier windows. The abbreviation “W” indicates the wild soybean; “C_” denoted the cultivated soybean; “L” represents the landraces; “I” denotes the improved cultivars. SR indicated the Chinese Southern region; CR implied the Chinese Central region surrounding the mid-down stream of Yellow River valley; NR stand for the Chinese Northern region plus Japan, Korean peninsula and Russian Far East region. “Am” pointed to America.

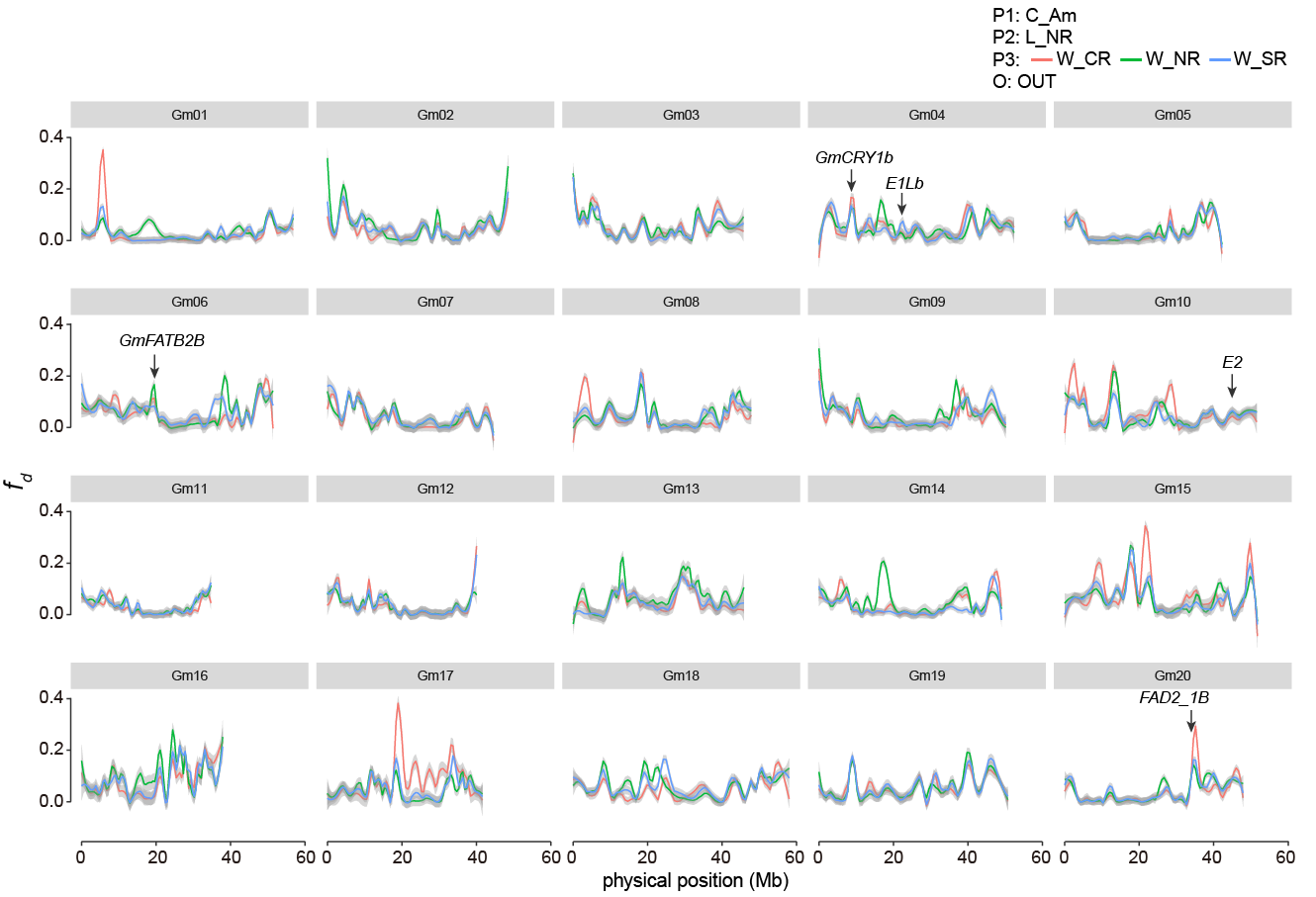

**Fig. S11.** **The distribution of *f*_d_ values in the landrace sub-population from the northern region (L_NR) with potential introgression from three wild sub-populations, respectively.** The arrows denoted some characterized genes located in the outlier windows. The abbreviation “W” indicates the wild soybean; “C_” denoted the cultivated soybean; “L” represents the landraces; “I” denotes the improved cultivars. SR indicated the Chinese Southern region; CR implied the Chinese Central region surrounding the mid-down stream of Yellow River valley; NR stand for the Chinese Northern region plus Japan, Korean peninsula and Russian Far East region. “Am” pointed to America.

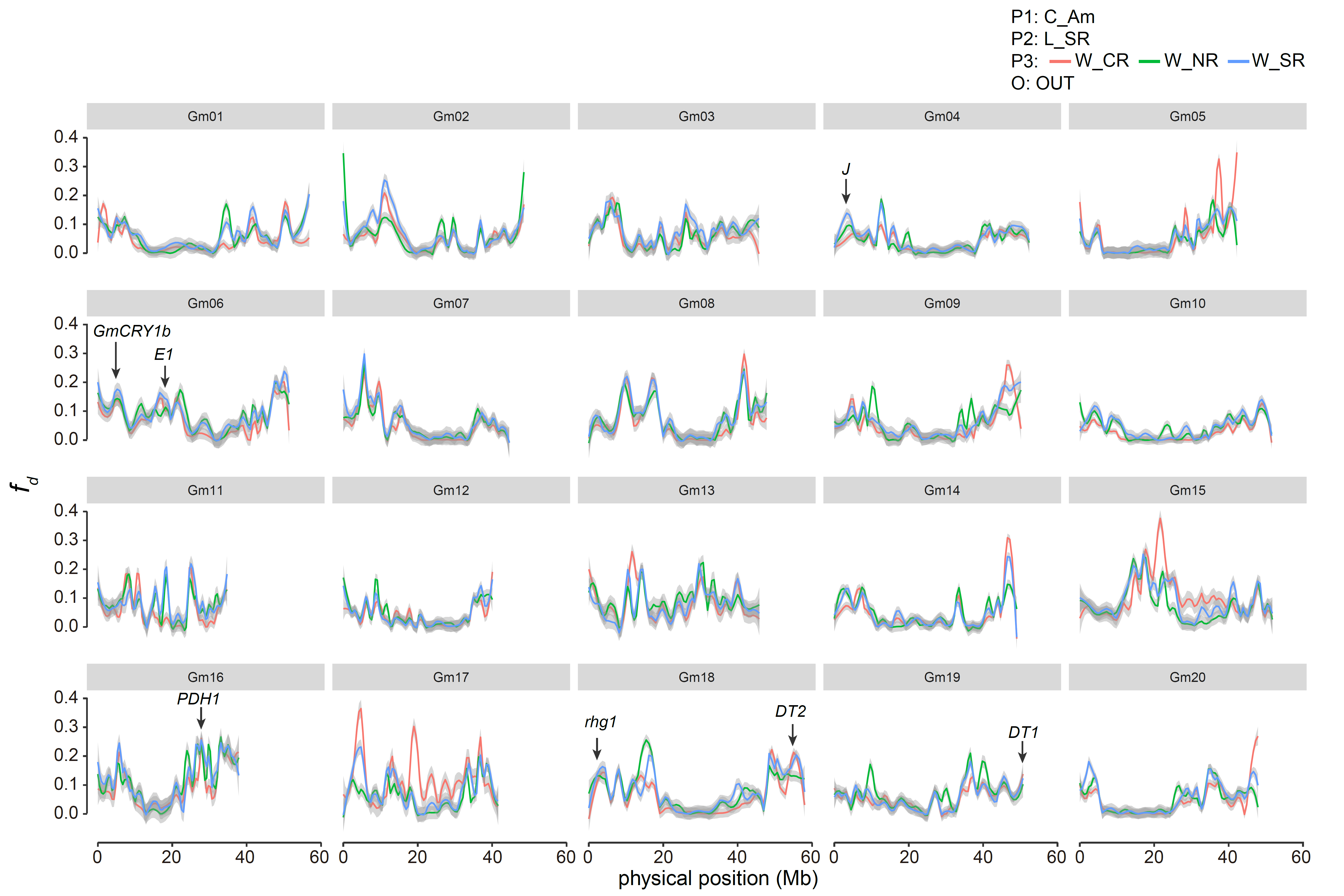

**Fig. S12.** **The distribution of *f*_d_ values in the landrace sub-population from the southern region (L_SR) with potential introgression from three wild sub-populations, respectively.** The arrows denoted some characterized genes located in the outlier windows. The abbreviation “W” indicates the wild soybean; “C_” denoted the cultivated soybean; “L” represents the landraces; “I” denotes the improved cultivars. SR indicated the Chinese Southern region; CR implied the Chinese Central region surrounding the mid-down stream of Yellow River valley; NR stand for the Chinese Northern region plus Japan, Korean peninsula and Russian Far East region. “Am” pointed to America.

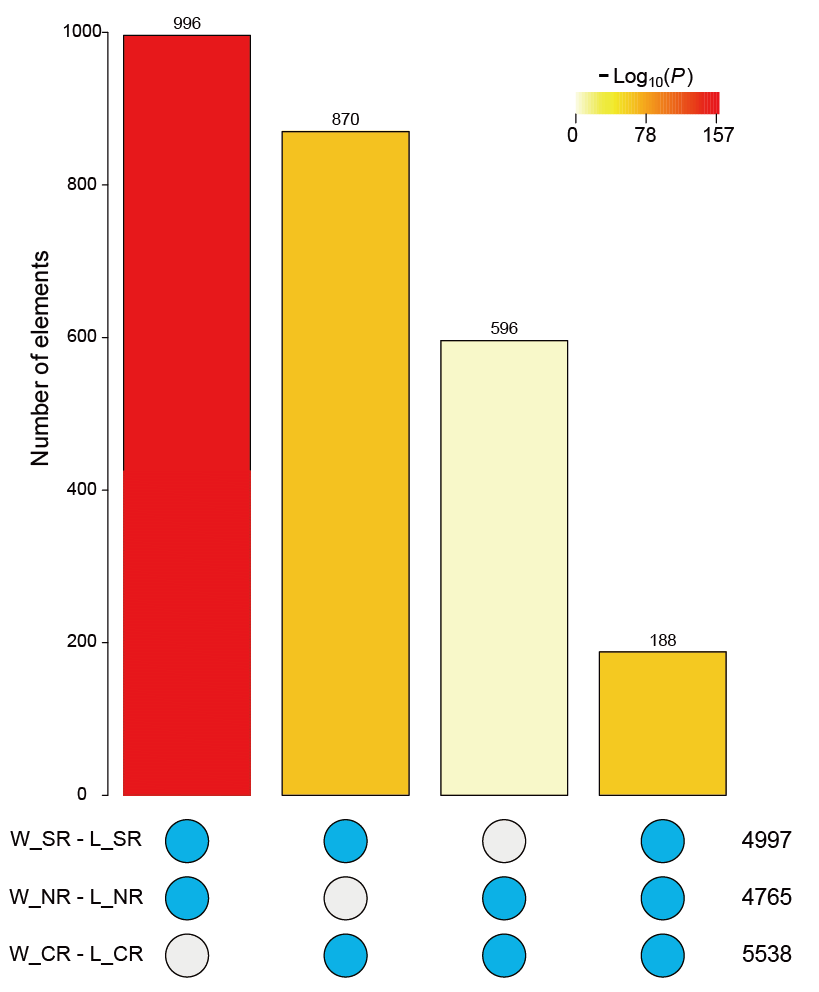

**Fig. S13. The sharing of *f*_d_ outlier windows among the three landrace sub-populations. The color of the bar indicates the significance level.** Abbreviations: “W_” represented the wild soybean and “L_” denoted the landrace; SR indicated the Chinese Southern region; CR implied the Chinese Central region surrounding the mid-down stream of Yellow River valley; NR stand for the Chinese Northern region plus Japan, Korean peninsula and Russian Far East region.

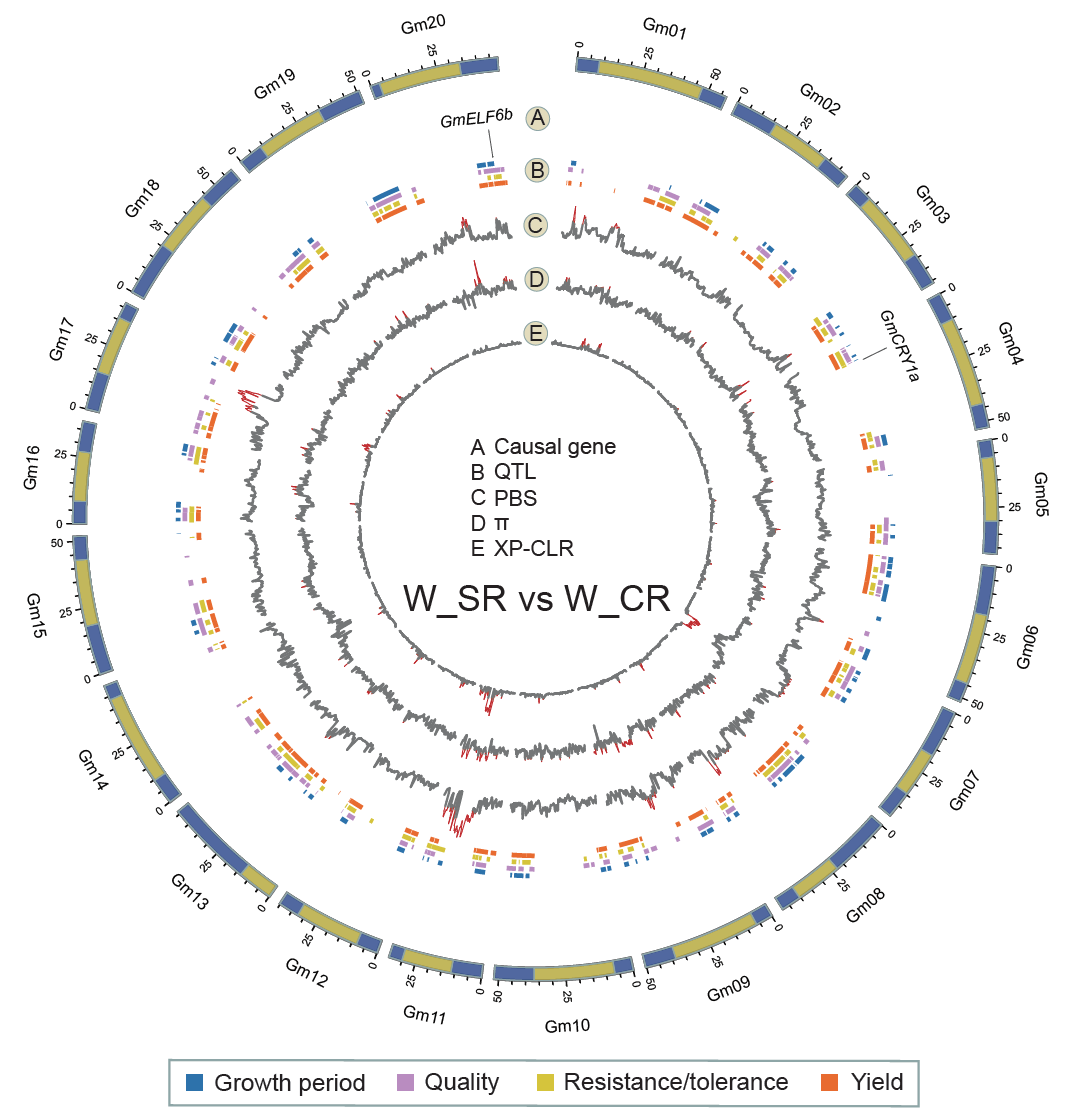

**Fig. S14. Genome-wide selection signatures in W_CR compared with W_SR on the basis of 8,785,134 SNPs.** The outermost ring indicated 20 chromosomes. Functional genes with selection signals were marked in ring A, and four functional QTLs were showed in ring B with different colors. Rings C to E represented selective signals for PBS, *θπ* ratio and XP-CLR, respectively. The genomic region with selection signal (the threshold value was set as Top 1%) were indicated in red.

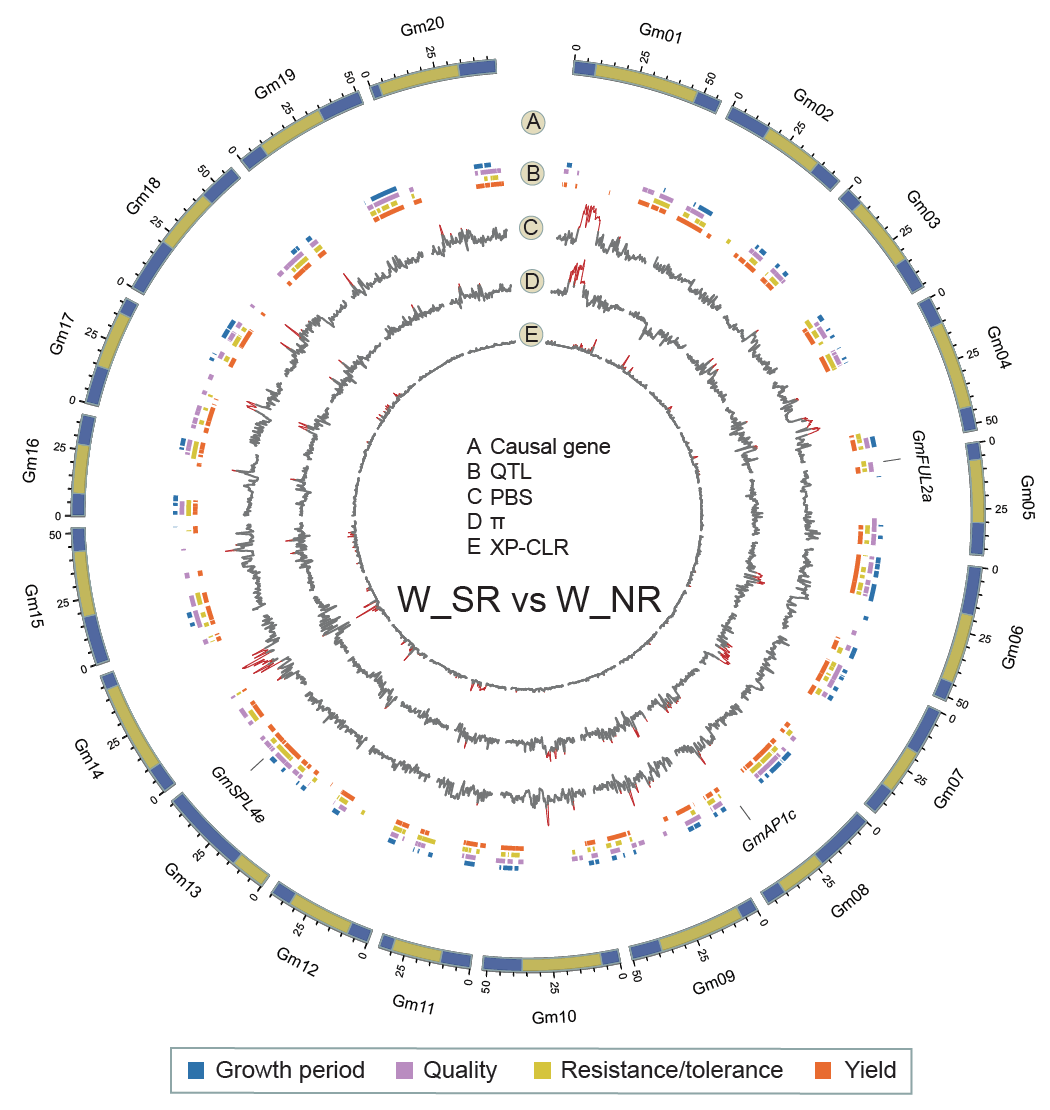

**Fig. S15. Genome-wide selection signatures in W_NR compared with W_SR on the basis of 8,785,134 SNPs.** The outermost ring indicated 20 chromosomes. Functional genes with selection signals were marked in ring A, and four functional QTLs were showed in ring B with different colors. Rings C to E represented selective signals for PBS, *θπ* ratio and XP-CLR, respectively. The genomic region with selection signal (the threshold value was set as Top 1%) were indicated in red.

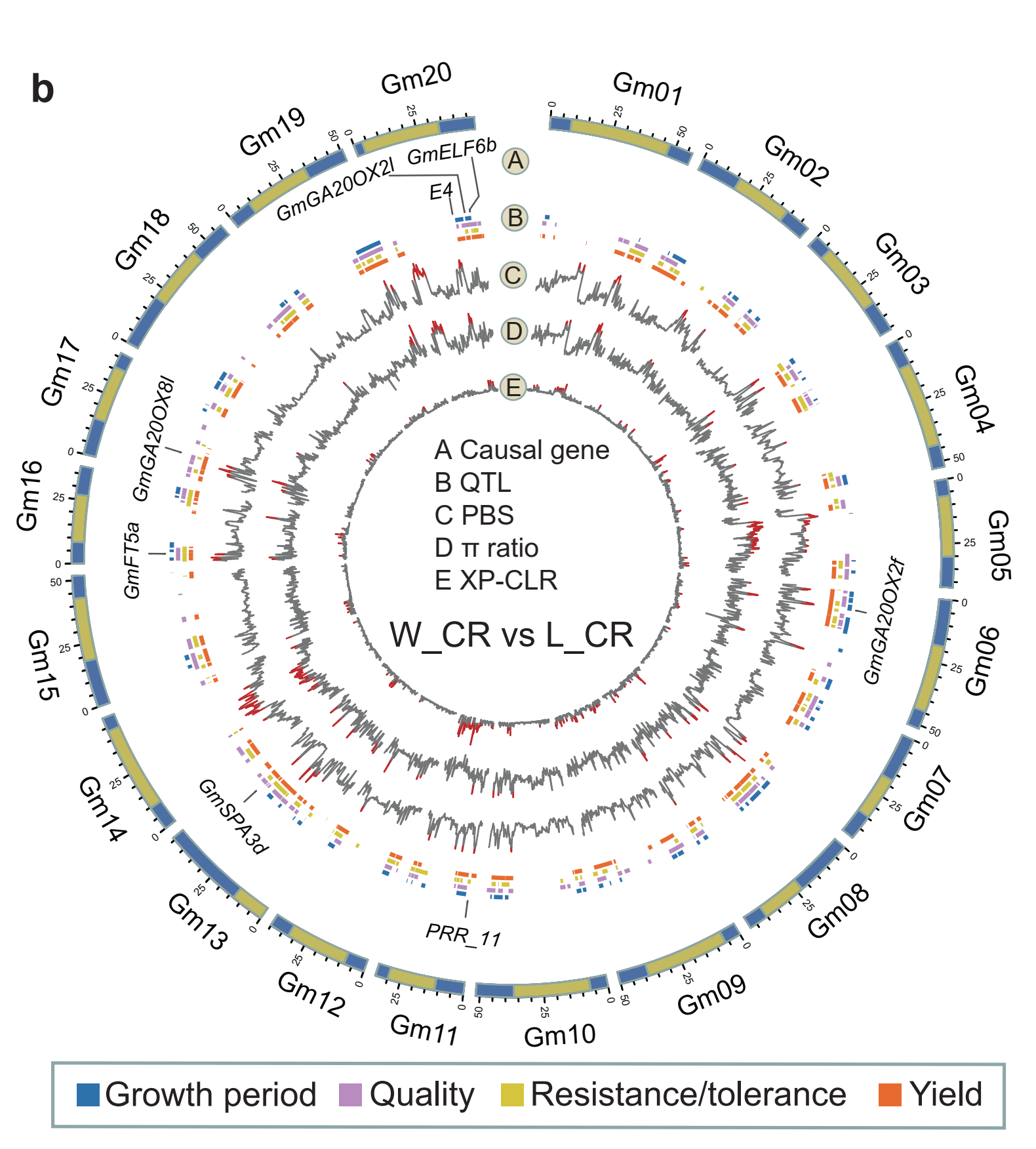

**Fig. S16. Genome-wide selection signatures of the domestication on the basis of 8,785,134 SNPs.** The outermost ring indicated 20 chromosomes. Functional genes with selection signals were marked in ring A, and four functional QTLs were showed in ring B with different colors. Rings C to E represented selective signals for PBS, *θπ* ratio and XP-CLR, respectively. The genomic region with selection signal (the threshold value was set as Top 1%) were indicated in red.

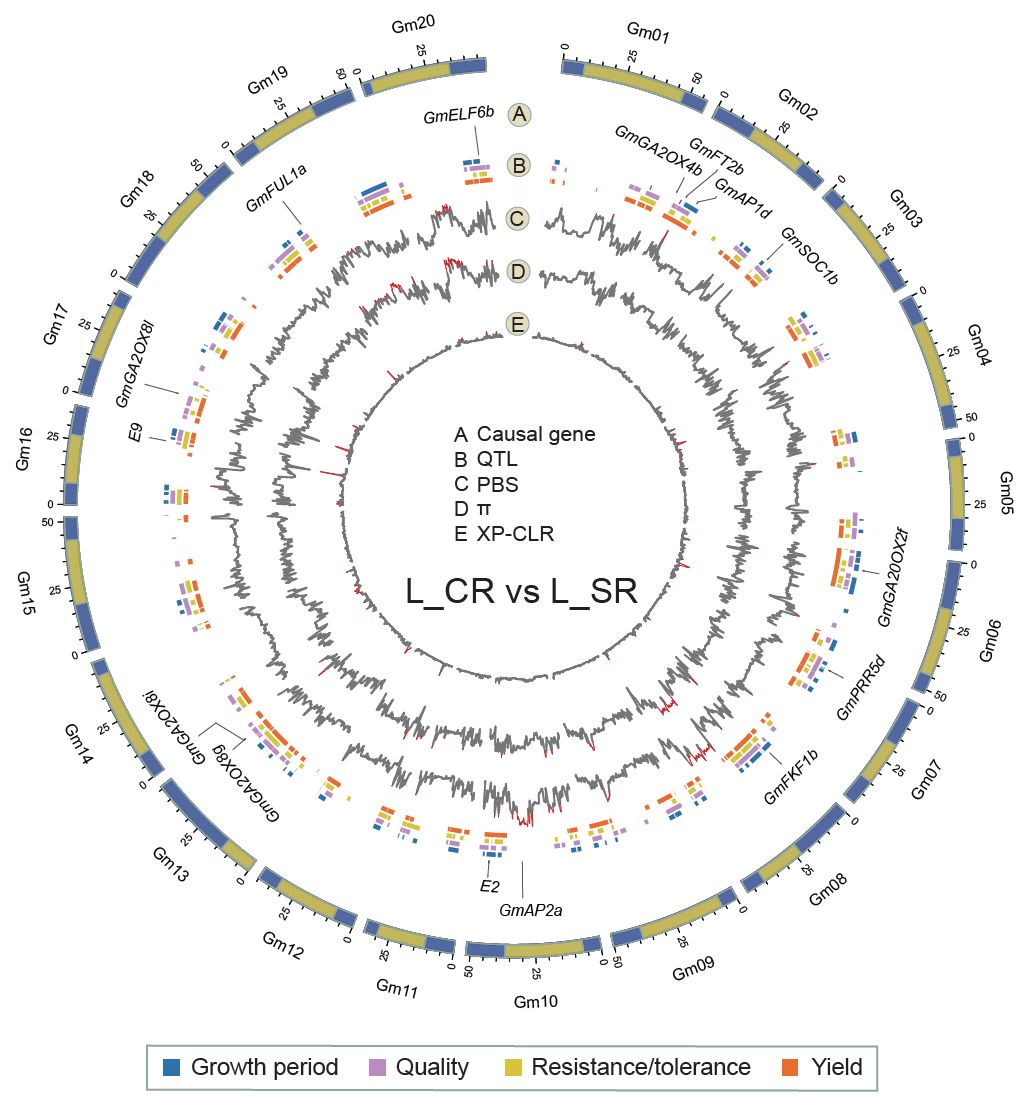

**Fig. S17. Genome-wide selection signatures in L_SR compared with L_CR on the basis of 8,785,134 SNPs.** The outermost ring indicated 20 chromosomes. Functional genes with selection signals were marked in ring A, and four functional QTLs were showed in ring B with different colors. Rings C to E represented selective signals for PBS, *θπ* ratio and XP-CLR, respectively. The genomic region with selection signal (the threshold value was set as Top 1%) were indicated in red.

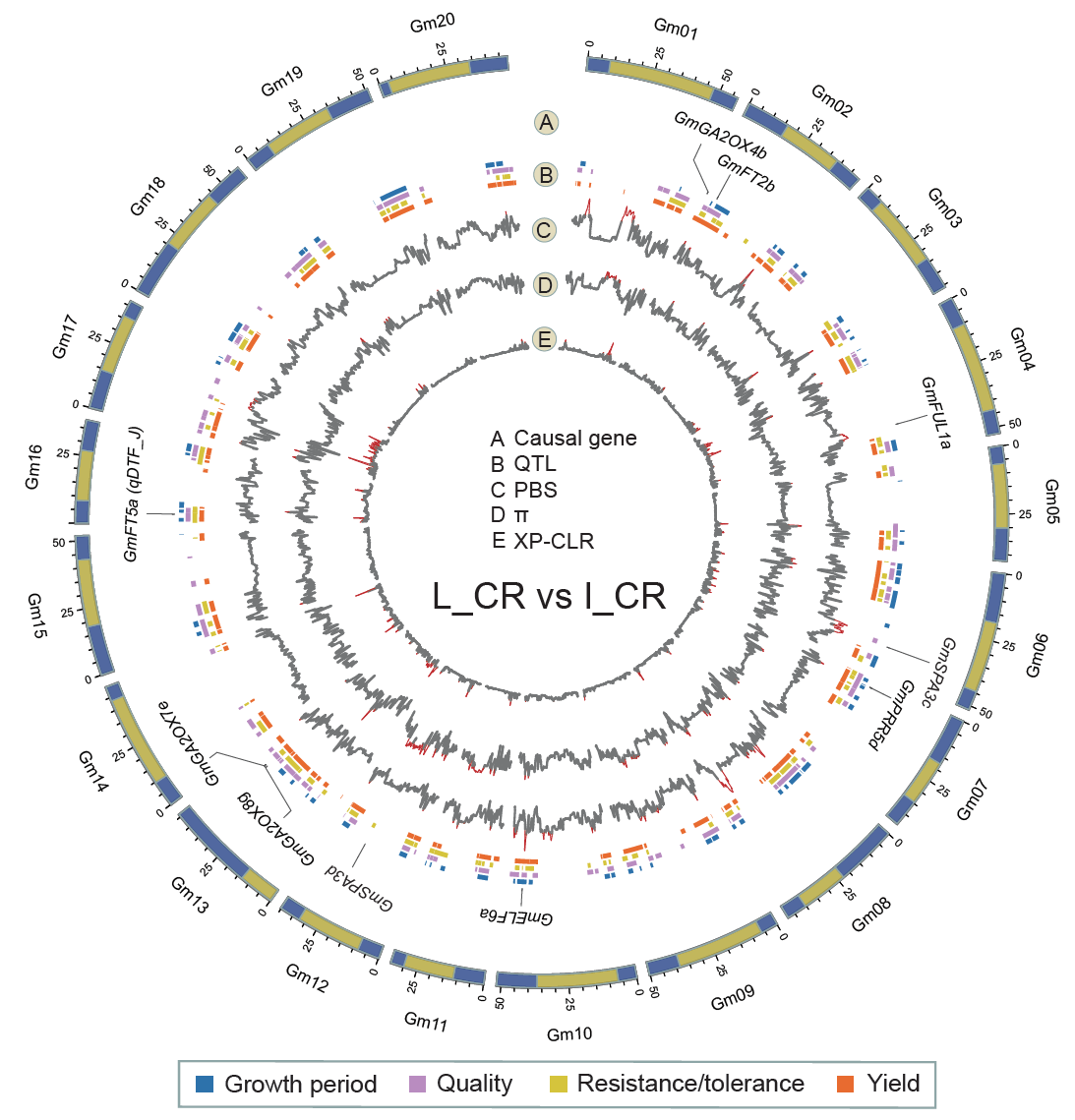

**Fig. S18. Genome-wide selection signatures in I_CR compared with L_CR on the basis of 8,785,134 SNPs.** The outermost ring indicated 20 chromosomes. Functional genes with selection signals were marked in ring A, and four functional QTLs were showed in ring B with different colors. Rings C to E represented selective signals for PBS, *θπ* ratio and XP-CLR, respectively. The genomic region with selection signal (the threshold value was set as Top 1%) were indicated in red.

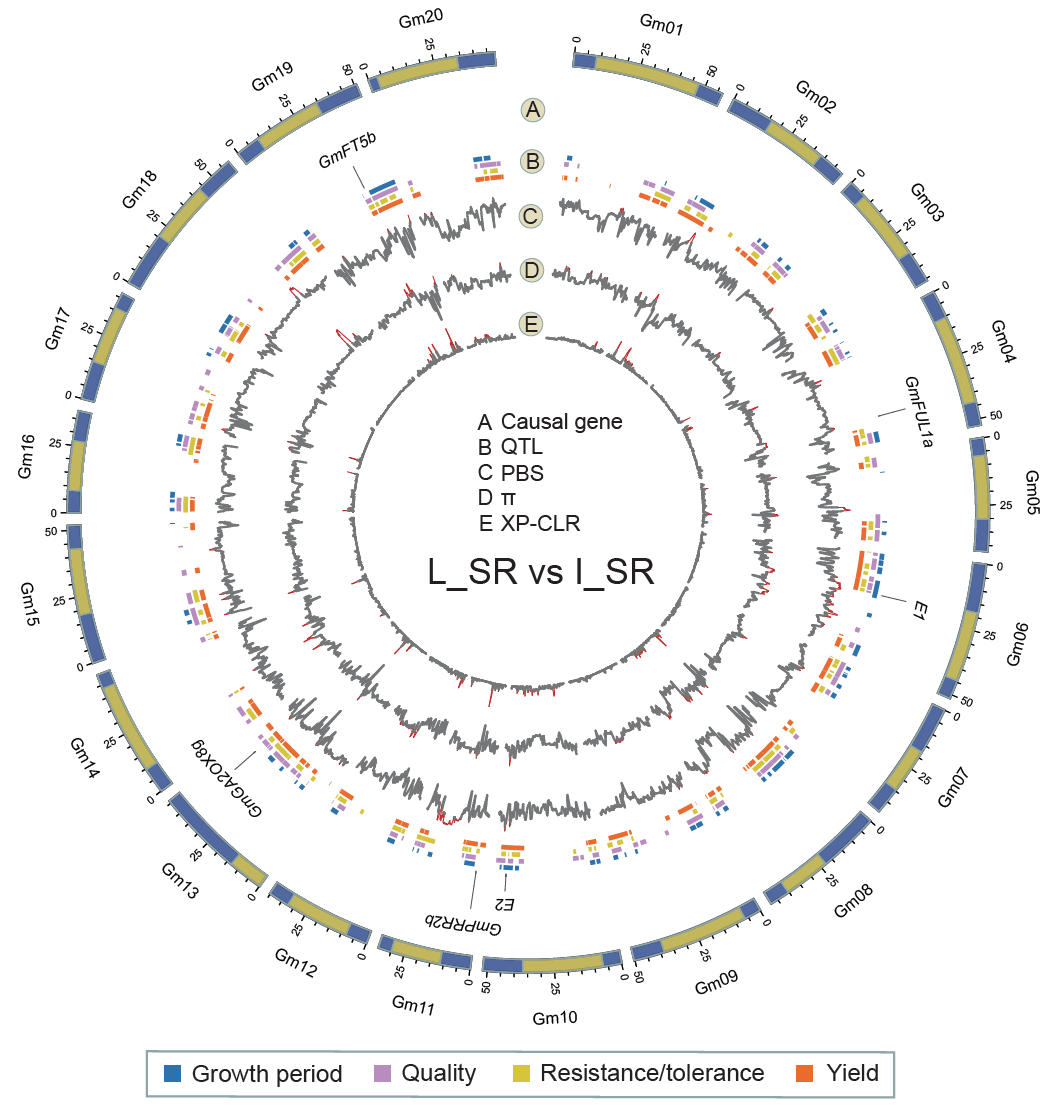

**Fig. S19. Genome-wide selection signatures in I_SR compared with L_SR on the basis of 8,785,134 SNPs.** The outermost ring indicated 20 chromosomes. Functional genes with selection signals were marked in ring A, and four functional QTLs were showed in ring B with different colors. Rings C to E represented selective signals for PBS, *θπ* ratio and XP-CLR, respectively. The genomic region with selection signal (the threshold value was set as Top 1%) were indicated in red.

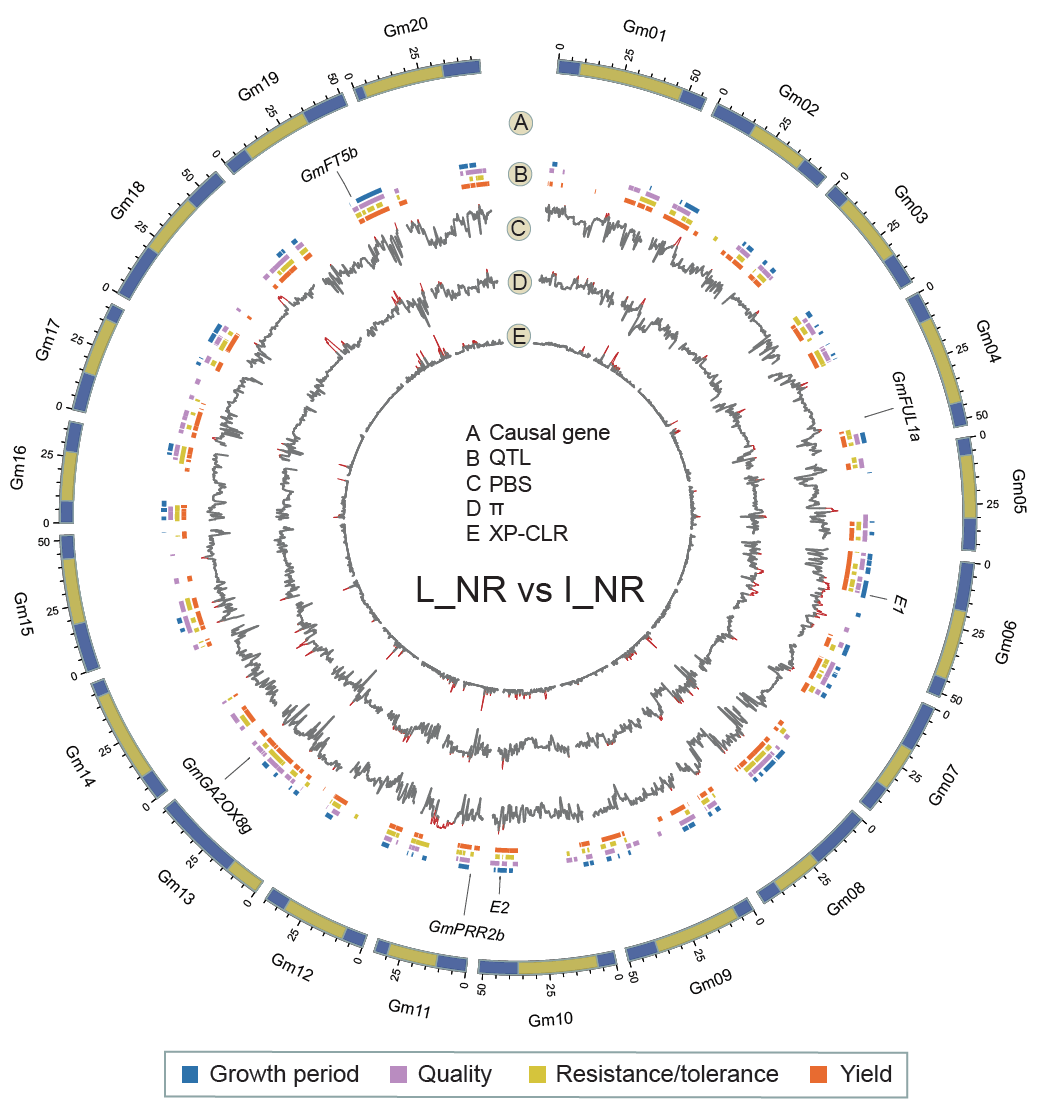

**Fig. S20. Genome-wide selection signatures in I_NR compared with L_NR on the basis of 8,785,134 SNPs.** The outermost ring indicated 20 chromosomes. Functional genes with selection signals were marked in ring A, and four functional QTLs were showed in ring B with different colors. Rings C to E represented selective signals for PBS, *θπ* ratio and XP-CLR, respectively. The genomic region with selection signal (the threshold value was set as Top 1%) were indicated in red.

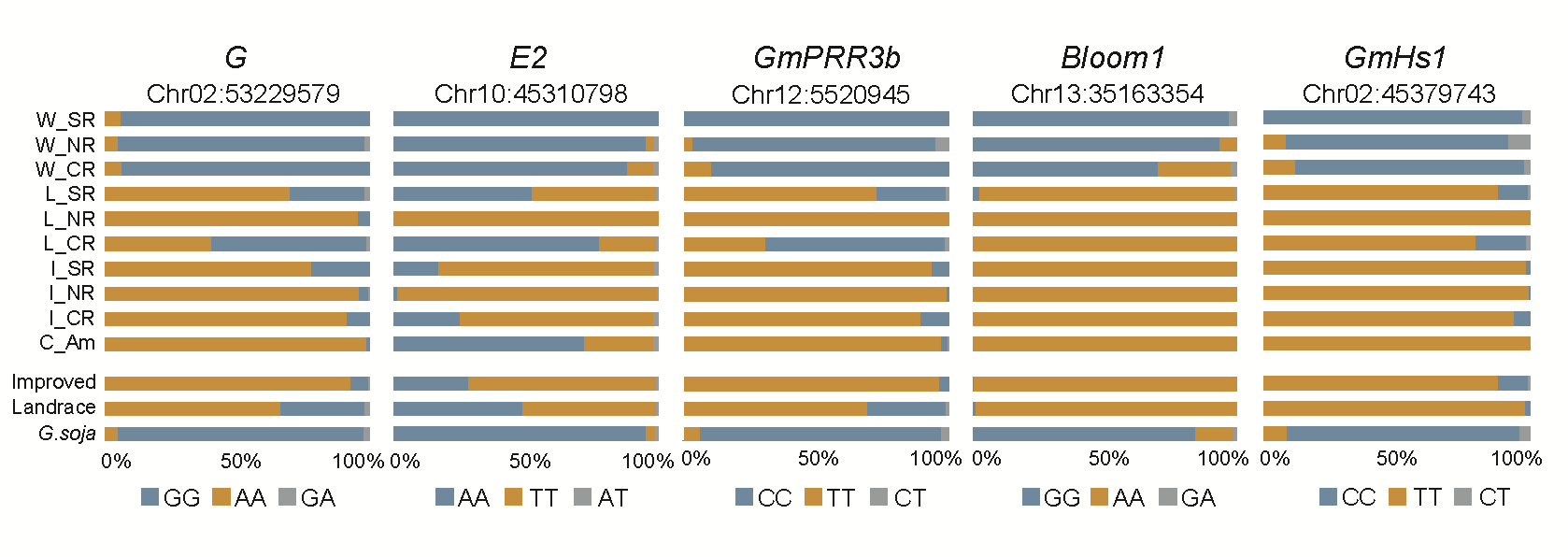

**Fig. S21. Allele frequencies of causal SNPs of five known genes with selection signals.**

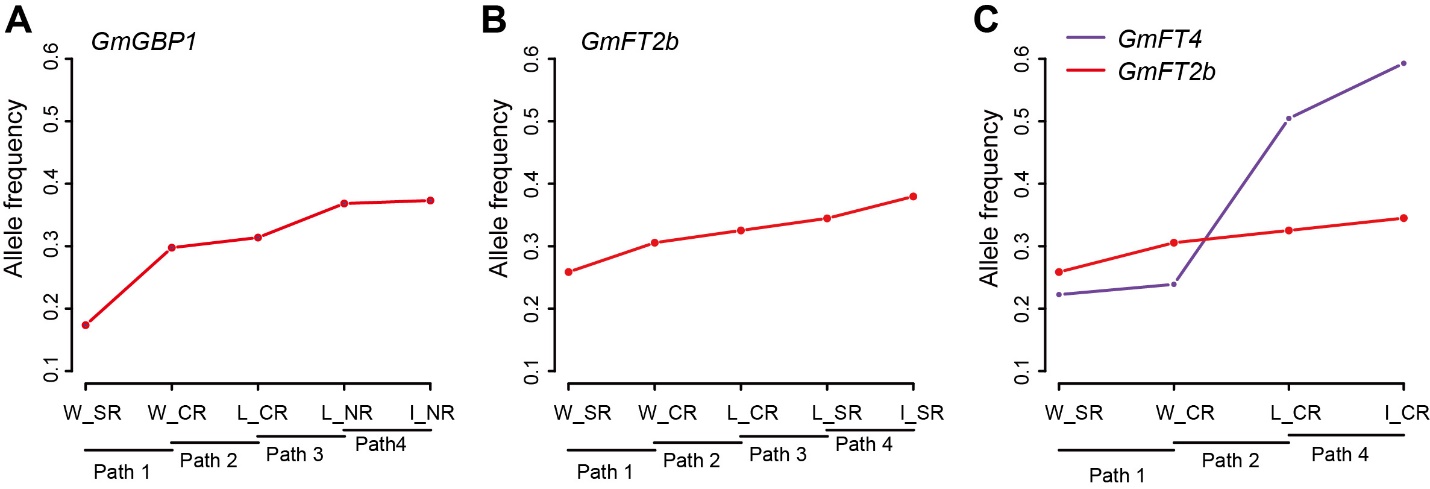

**Fig. S22. Plot of allele frequencies changes of three flowering cloned genes across integrating putative intact evolutionary stages in Northern (A), Southern (B), and Central regions (C).**

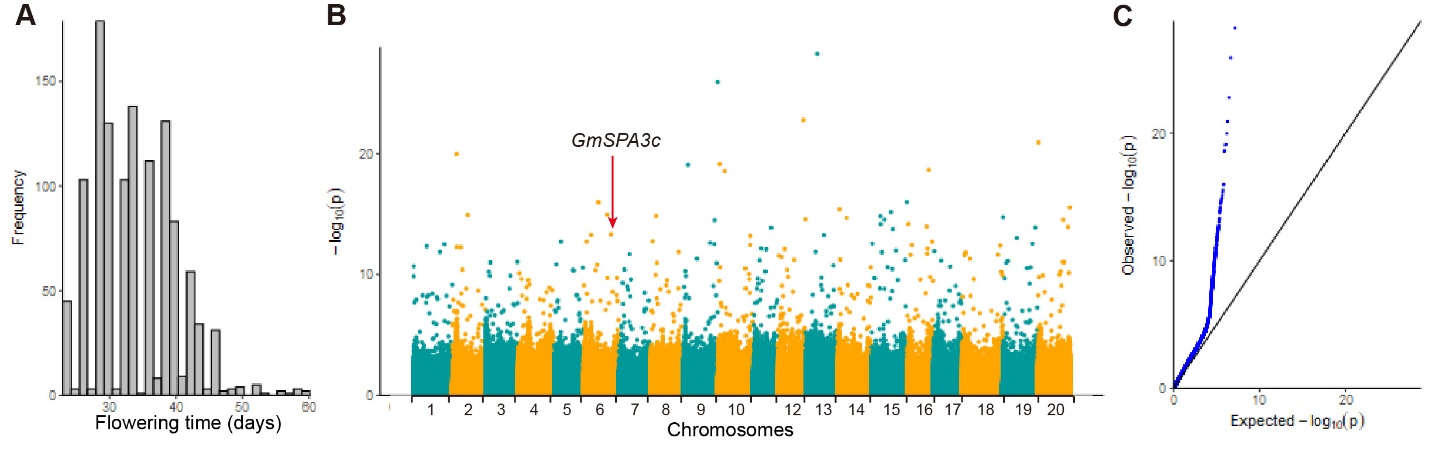

**Fig. S23.** **GWAS of flowering phenotypic trait in 2,214 soybeans using FarmCPU,** including frequency distribution of phenotypic data (**A**), Manhattan plot (**B**) and quantile-quantile plot (**C**). The horizontal dash line indicates the significant threshold (1×10^-6^).

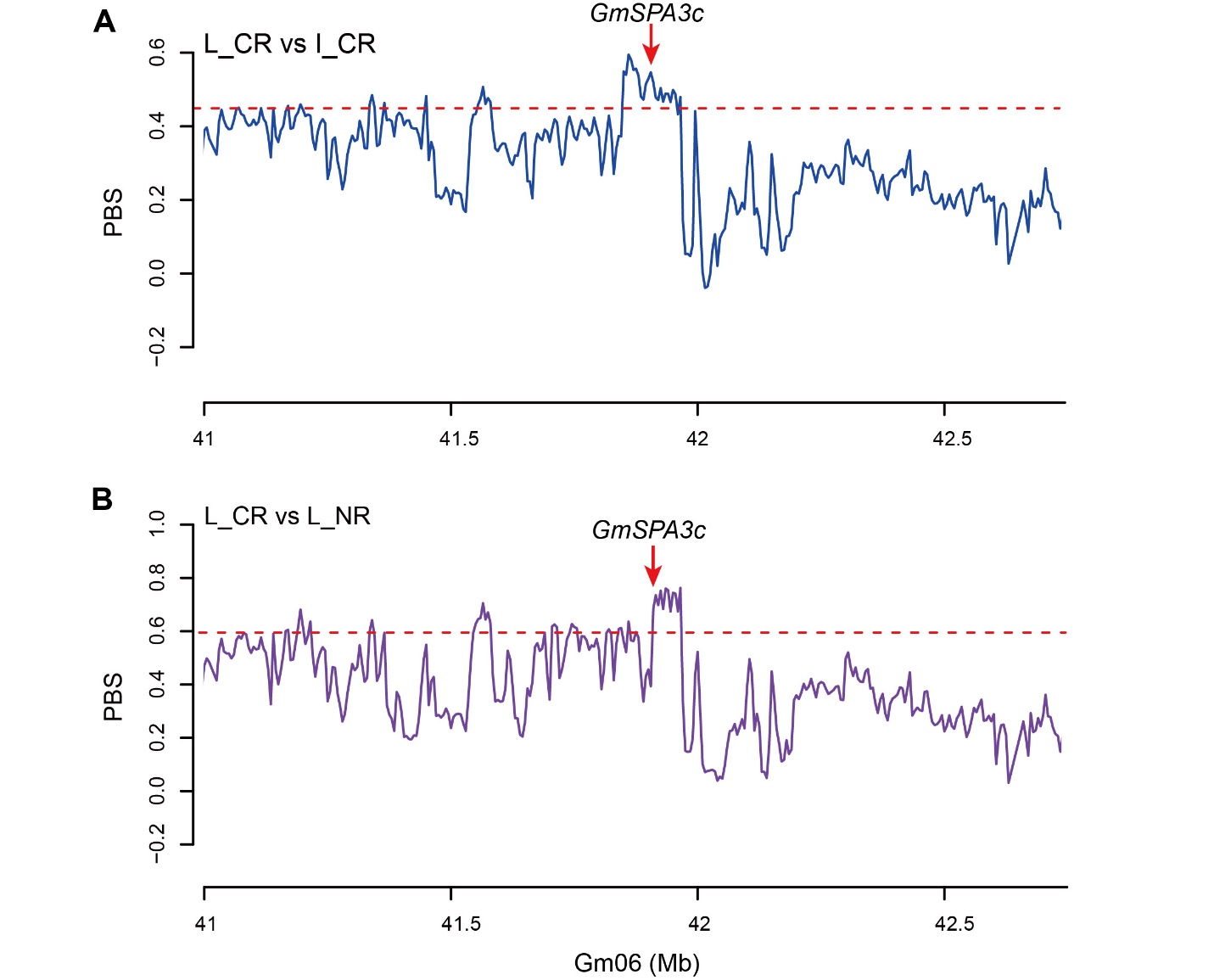

**Fig. S24. Selective sweeps of *GmSPA3c* identified with the PBS statistic.**

(**A**) selective signals of *GmSPA3c* in I_CR compared with L_CR. (**B**) selective signals of *GmSPA3c* in I_NR compared with L_NR. Black dotted line indicated the threshold at the top 5% PBS value.

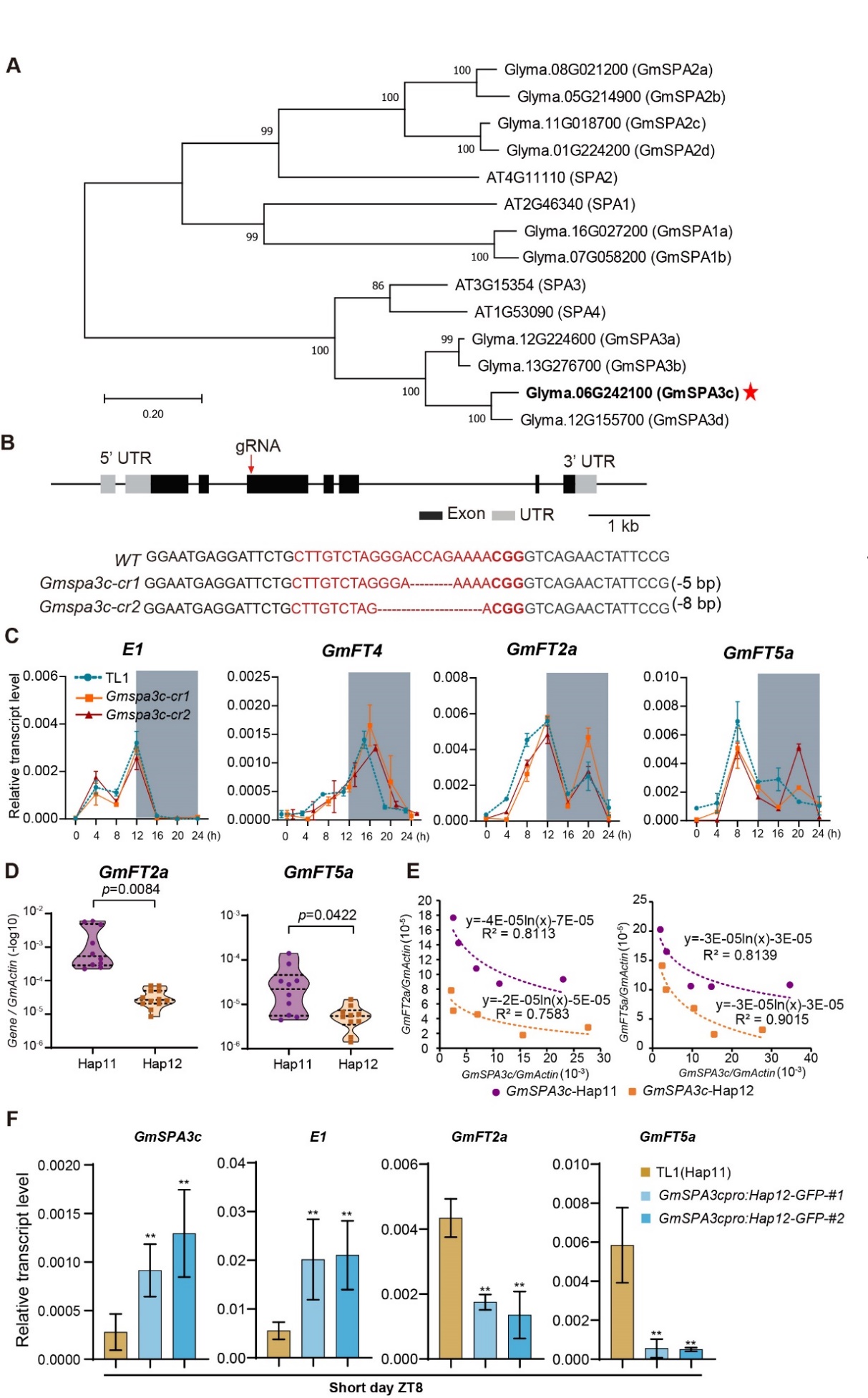

**Fig.S25. The phylogenetic and functional analysis of the *GmSPA3c* gene.**

(**A**) Phylogenetic tree of SPA proteins from Arabidopsis and soybean using neighbor-joining method with the MEGA5 software. (**B**) Guided RNAs (sgRNA, red arrows) were designed to target the third exon of *GmSPA3c* Hap11. The mutant sequences of two representative homozygous mutants (*Gmspa3c*-*cr1* and *Gmspa3c*-*cr2*) at T2 generation are shown. The target sites of gRNA were highlighted in red letters with the protospacer-adjacent motif (PAM) in bold. The red dashed lines within the target sites denote nucleotide deletions. (**C**) The dynamic transcriptional level of *E1, GmFT4, GmFT2a,* and *GmFT5a* in the *Gmspa3c* mutants and TL1 under short day conditions. Mean values ± s.d. (n = 3) are shown. (**D**) The transcriptional levels of *GmFT2a* and *GmFT5a* at peaking points in different accessions carrying Hap11 or Hap12. Details about the accessions were provided in Supplementary Table 8. The plants were grown under long day conditions for 21 days, then the second trifoliate leaves were collected at ZT4 for *GmFT2a* and ZT8 for *GmFT5a* respectively. The comparisons were performed using student’s *t*-test (n > 8). (**E**) Correlation analysis between *GmSPA3c* (Hap11 and Hap12) and *GmFT2a/5a* mRNA levels in calluses transformed with 35s droved *GmSPA3c* (Hap11 and Hap12) at ZT4. (**F**) The transcriptional levels of *GmSPA3c, E1, GmFT2a,* and *GmFT5a* in the indicated lines at ZT8 under short day conditions.
